# Immune targets for schistosomiasis control identified by a genome-wide association study of East African snail vectors

**DOI:** 10.1101/2024.08.30.610565

**Authors:** Tom Pennance, Jacob A Tennessen, Johannie M Spaan, Tammie McQuistan, George Ogara, Fredrick Rawago, Kennedy Andiego, Boaz Mulonga, Meredith Odhiambo, Martin W Mutuku, Gerald M Mkoji, Eric S Loker, Maurice R Odiere, Michelle L Steinauer

**Affiliations:** College of Osteopathic Medicine of the Pacific – Northwest, Western University of Health Sciences, Lebanon, OR, USA; Harvard T.H. Chan School of Public Health, Boston, MA, USA; Centre for Global Health Research, Kenya Medical Research Institute (KEMRI), P. O. Box 1578- 40100, Kisumu, Kenya; Centre for Biotechnology Research and Development, Kenya Medical Research Institute (KEMRI), P.O. Box 54840–00200, Nairobi, Kenya; Department of Biology, Center for Evolutionary and Theoretical Immunology, Parasite Division Museum of Southwestern Biology, University of New Mexico, Albuquerque, New Mexico, USA

**Keywords:** *Schistosoma*, *Biomphalaria*, GWAS, amplicon panel, pathogen resistance

## Abstract

Schistosomiasis, afflicting >260 million people worldwide, could be controlled by preventing infection of freshwater snail vectors. Intestinal schistosomiasis, caused by *Schistosoma mansoni*, occurs predominantly in Sub-Saharan Africa and is vectored by *Biomphalaria sudanica* and related *Biomphalaria* species. Despite their importance in transmission, very little genomic work has been initiated in African snails, thus hindering development of novel control strategies. To identify genetic factors influencing snail resistance to schistosomes, we performed a pooled genome-wide association study (pooled-GWAS) on the offspring of *B. sudanica* collected from a persistent hotspot of schistosomiasis in Lake Victoria, Kenya, and exposed to sympatric *S. mansoni*. Results of the pooled-GWAS were used to develop an amplicon panel to validate candidate loci by genotyping individual snails. This validation revealed two previously uncharacterized, evolutionarily dynamic regions, *SudRes1* and *SudRes2*, that were significantly associated with resistance. *SudRes1* includes receptor-like protein tyrosine phosphatases and *SudRes2* includes a class of leucine-rich repeat-containing G-protein coupled receptors, both comprising diverse extracellular binding domains, suggesting roles in pathogen recognition. No loci previously tied to schistosome resistance in other snail species showed any association with compatibility suggesting that loci involved in the resistance of African vectors differ from those of neotropical vectors. Beyond these two loci, snail ancestry was strongly correlated with schistosome compatibility, indicating the importance of population structure on transmission dynamics and infection risk. These results provide the first detail of the innate immune system of the major schistosome vector, *B. sudanica*, informing future studies aimed at predicting and manipulating vector competence.

**Significance Statement:** Although schistosomiasis-associated morbidity and mortality have reduced significantly due to chemotherapy, interruption of transmission, a WHO goal, requires complimentary novel snail vector-focused interventions. We performed a genome-wide association study of snails exposed to schistosomes in an endemic area of high transmission in Kenya. We identified two snail genomic regions that were associated with snail immunity to schistosomes, and which had not previously been tied to parasite infection. The characterized protein structures are plausibly consistent with a role in host-parasite interaction. Therefore, they are new, potential targets for schistosomiasis control. These results show the need for focused research on transmission-relevant vectors and their genetic variants that could have a large impact on schistosome transmission dynamics and human health risk.

## Main Text

Schistosomiasis is a global scourge, taking a large toll on people who have the fewest resources. Affecting over 260 million people, it is the parasitic disease with the greatest impact on health worldwide after malaria (1, 2). Within the last decade, schistosomiasis control program goals have shifted from reduction of morbidity to elimination or interruption of schistosomiasis as a public health problem by 2030 (1, 3, 4). However, the toolbox with which to combat schistosome transmission has remained virtually the same, dominated by one main approach: mass drug administration (MDA) of praziquantel (5). It is increasingly recognized that in addition to chemotherapy, successful control and elimination will require targeting the aquatic snails which serve as intermediate hosts of the schistosome parasites and transmit them to humans (6, 7). Part of the reason MDA alone is insufficient is that even with effective drug treatment, people become rapidly reinfected by infected snails in the environment (8–10). Schistosomes form chronic infections in snails and continually release hundreds to thousands of infectious stages (cercariae) into the environment daily (11).

Historically, schistosomiasis control programs that are focused on snail control have been the most successful at reducing or eliminating schistosomiasis (12, 13); however, snail-directed control methods are limited and have negative impacts. Molluscicides are indiscriminately toxic and are impractical to apply to vast habitats (12, 14). Furthermore, snail population rebound post-molluscicide application is predicted to increase schistosome transmission (15).

Given the absence of suitable snail vector control methods for contemporary public health interventions, there is a need for new approaches to be developed (16). Genomic and transcriptomic data and resources enable approaches like genome-wide association studies (GWAS) that identify genomic regions of snail vectors involved in resisting schistosome infection (17). Once identified, snail genes and genomic variants associated with resistance to schistosomes could be monitored in wild populations and potentially be manipulated so that snail resistance to schistosomes is enhanced and transmission to humans is interrupted.

The feasibility of manipulating snail resistance to schistosomes may follow similar approaches to engineering resistant hosts for disease control using CRISPR-Cas and associated gene drive technologies (18, 19). However, much of the elegant work regarding transcriptomics and genomics of schistosome-snail compatibility has only addressed these questions in laboratory models of the South American vector of *Schistosoma mansoni, Biomphalaria glabrata* (17, 20–24). Very little is known regarding how this body of knowledge will translate to African vectors of *S. mansoni*, through which 90% of *S. mansoni* transmission occurs (25). The recent publication of two African *Biomphalaria* species genomes and transcriptomes (26, 27) provide a path toward molecular-informed snail control in hotspots of transmission.

With the goal of identifying and describing the genetic architecture underlying resistance of African snails to *S. mansoni*, we performed a pooled genome-wide association study (pooled-GWAS) (28) using a wild population of *Biomphalaria sudanica* originating from the shores of Lake Victoria closest to a persistent hotspot of schistosomiasis in Kanyibok, western Kenya (10). We identify two large-effect loci and a strong influence of ancestry on snail resistance to schistosome infection, defined here as complete parasite clearance (including parasite DNA) from snail tissue following exposure. These results reveal how immunogenetics and population demographics contribute to vectorial competence in a natural vector population with direct impact on human health.

## Results

### Pooled-GWAS reveals multiple variants strongly enriched in resistant snails

Of 1400 F1 *B. sudanica*, whose parents originated from Anyanga Beach (Lake Victoria, western Kenya), exposed to eight freshly hatched *S. mansoni* miracidia from local schoolchildren, 1,109 snails remained in the GWAS study after excluding 254 that died prior to screening for infection and 37 that yielded insufficient genomic DNA (gDNA) quality. The final sample set comprised 615 and 393 snails that were positive (i.e. releasing *S. mansoni* cercariae) or negative (i.e. not releasing *S. mansoni* cercariae nor PCR positive (29)), respectively. Snails that were negative for cercariae but positive for *S. mansoni* gDNA were not considered further. Two equal mass gDNA pools for the pooled-GWAS comprised 493 positive and 295 negative snails. The pooled-GWAS sequencing (Illumina paired-end 150 bp, NovaSeq 6000 S4 flow cell) yielded on average 1.5x coverage per snail (Dataset S1). A total of 4,498,972 variants were retained for analysis. Correlation between sequencing technical replicates of positive and negative pooled gDNA was weak but significantly positive, with 8-fold enrichment for outliers in the top 1% of both replicates (Fig. S1).

In the pooled-GWAS results, genotype-phenotype association *p* values ranged as low as 1e-30, including 45 variants (0.001%) with *p* ≤ 1e-15 and 1930 variants (0.04%) with *p* ≤ 2.5e-9 (Fig. 1). Rather than simply defining a genome-wide significance threshold to identify candidates to be validated, we prioritized genomic regions meeting a dual-variant criterion, whereby two or more proximate (<50 kb and >1.5 kb apart to ensure support from distinct read pairs) variants are strongly associated with *S. mansoni* resistance (arbitrary threshold of *p* ≤ 2.5e-9); such sliding dual-variant 50 kb windows encompass 18.625 Mb (2%) of the *B. sudanica* genome and contain 888 (46%) of variants with *p* ≤ 2.5e-9.

**Fig. 1.**
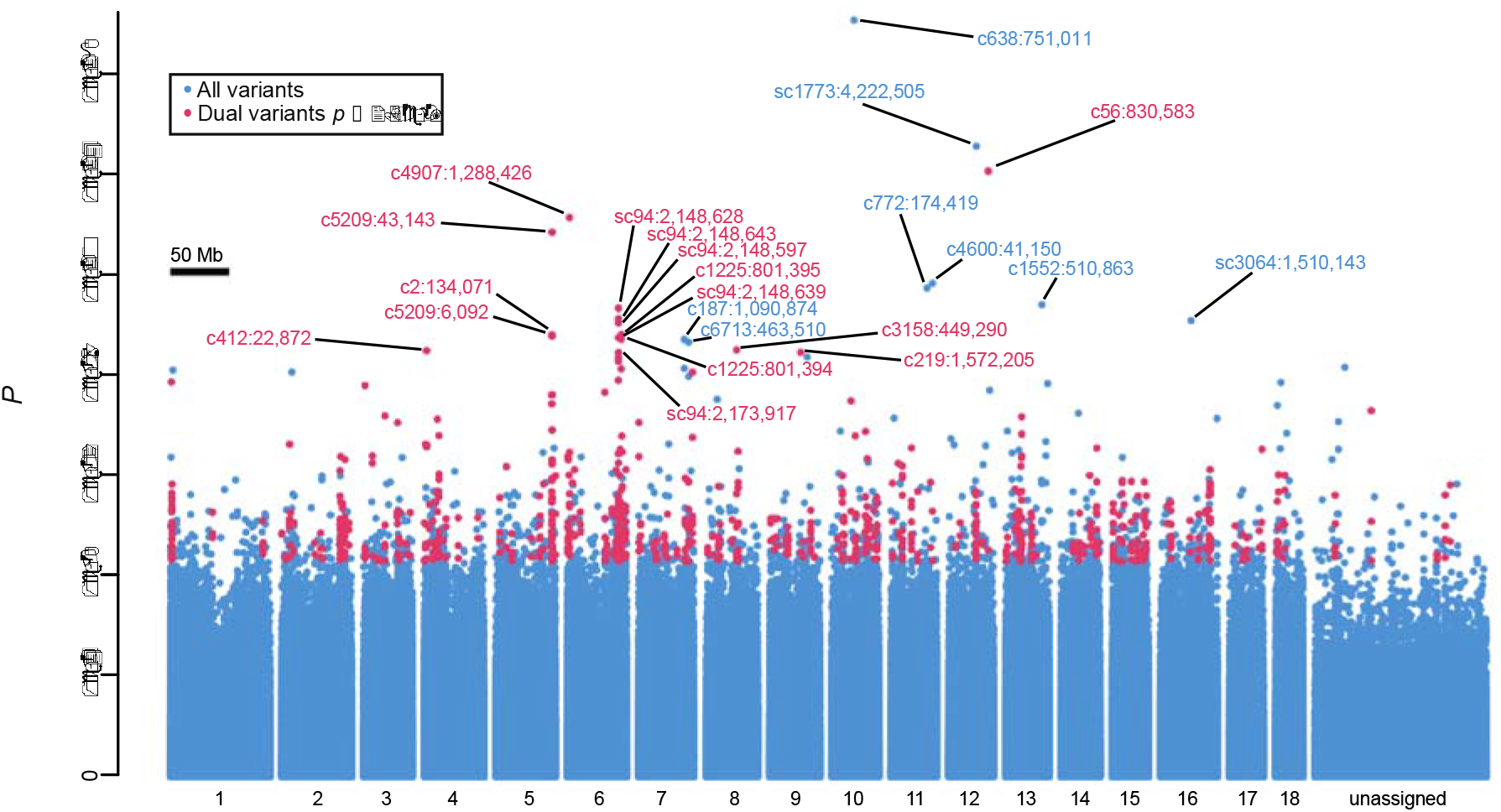
Results of the pooled genome-wide association study (pooled-GWAS) identifying variants associated with resistance to *Schistosoma mansoni* in the *Biomphalaria sudanica* genome. Fisher’s exact test *p* values for all pooled-GWAS variants, are arranged horizontally based on contig orthology to 18 chromosomes (x-axis labels) of the *B. glabrata* genome (xgBioGlab47.1, NCBI RefSeq: GCF_947242115.1) and linkage map analysis (Dataset S2, Fig. S2-S4). Pooled-GWAS dual-variants, defined as two or more proximate (<50 kb and >1.5 kb apart) variants strongly associated with *S. mansoni* resistance (*p* ≤ 2.5e-9) are red. All others are blue. All dual-variants and singleton-variants with p ≤ 1e-17 are labeled red and blue, respectively, with their contig and contig position. Unassigned contigs could not be unambiguously mapped and are mostly small and/or repetitive.

### Amplicon panel genotyping reveals population structure and validates variants associated with resistance

A multiplex amplicon panel was designed using the Genotyping-in-Thousands by sequencing method (30) to genotype variants in individual snails at 234 dual-variants and 12 singleton-variants with *p* < 1e-13 identified from the pooled-GWAS analysis (Dataset S3). The amplicon panel also contained 22 markers for *a priori* gene candidates and 201 ‘neutral’ markers to facilitate a linkage map for improved *B. sudanica* genome assembly (Dataset S2).

An independent set of 122 positive and 98 negative snails not included in the pooled-GWAS were reserved for validation of genomic variants via genotyping with the amplicon panel and are henceforth referred to as genotyped-validation snails. These independent genotyped-validation snails were used to validate the differential allele frequencies associated with *S. mansoni* resistance observed in the pooled-GWAS sequencing data. These were combined with a subset of the pooled-GWAS snails (genotyped-pooled-GWAS snails: 138 positive, 138 negative) for more precise estimates of ancestry and genotype frequencies.

The amplicon panel data (of which median missing data was 2% per locus and 2% per individual) revealed a signal of population structure, evident in both PCA (Fig. S5) and ADMIXTURE (31) analysis (Fig. 2A). With K=2 ancestral populations (CV = 0.53, versus 0.61 for K=1), ancestry from Population 2 ranged continuously from 0 to 1 and was significantly correlated with resistance (*p* < 1e-14). Mean Population 2 ancestry was 39% for positives and 62% for negatives. Snails were designated into two groups based on the percentage of their ancestry: 41% of snails had mostly Population 1 ancestry and were deemed “Group A”, while the remainder, with mostly Population 2 ancestry, were deemed “Group B”. Of the snails in Group A, 68% were positive. In Group B, 41% were positive. Thus, any marker differing in frequency between these ancestral populations could be correlated with infection phenotype, even if not linked to an etiological variant, and therefore could explain some outliers identified by the pooled-GWAS. Using markers diagnostic of both groups A and B, we found that *B. choanomphala*, a closely related deep-water taxon/eco-phenotype of *B. sudanica* also collected in Lake Victoria (32), and *B. sudanica* inbred lines originating from Lake Victoria (27), share ancestry with Group A, and thus the GWAS ancestry signal is not caused by interspecies hybridization (Fig. S6). The inferred percentage of Population 1 or 2 ancestry for each snail was similar when estimated using linkage map markers alone, and thus not driven by GWAS outliers (Fig. S7).

**Fig. 2.**
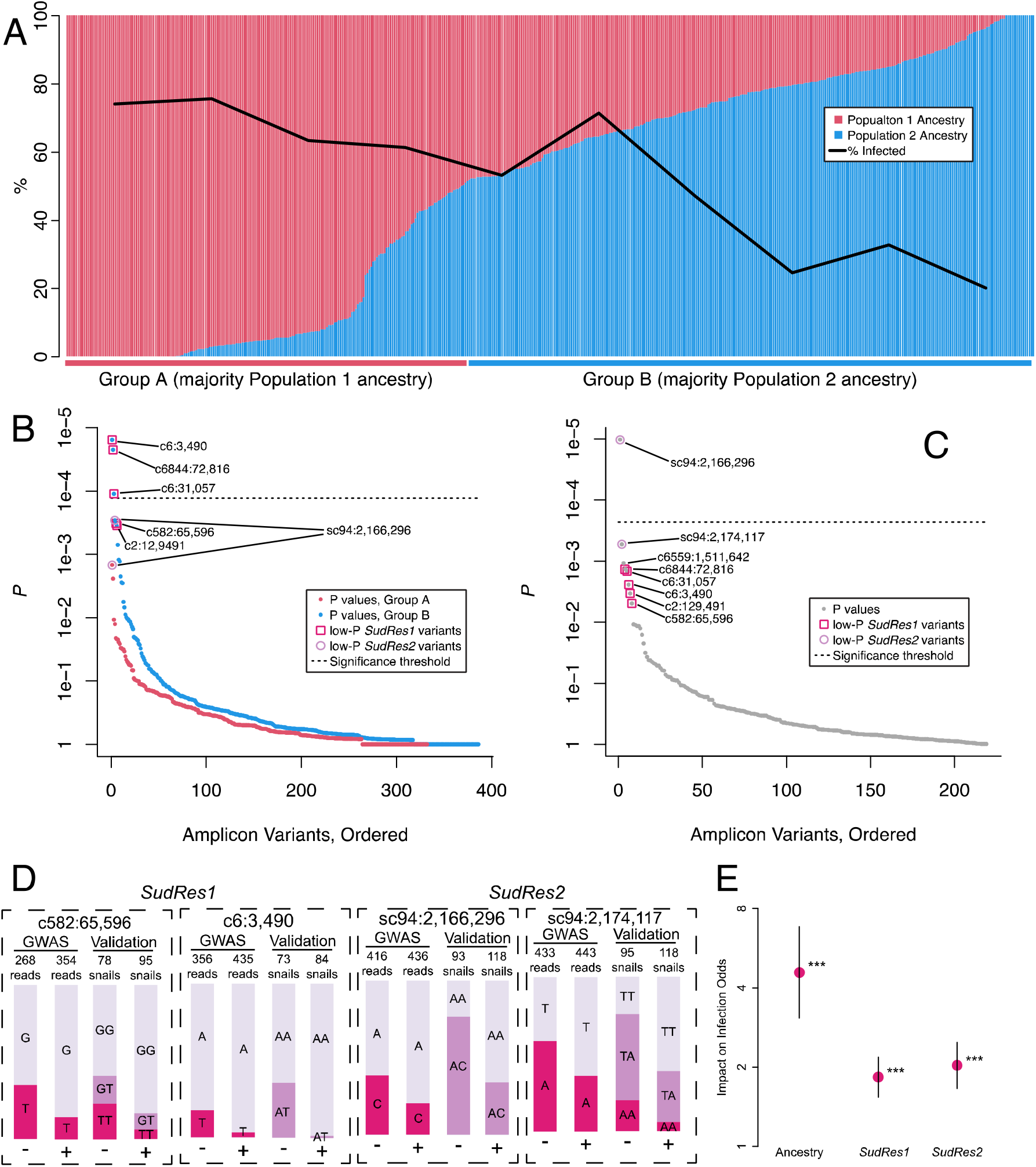
Amplicon panel validation of pooled-GWAS results. (A) Proportion of ancestry from *Biomphalaria sudanica* Population 1 (red) or Population 2 (blue) as predicted by ADMIXTURE (31) is correlated with *Schistosoma mansoni* infection (*p* < 1e-14), such that majority-Population 1 snails (Group A) are 68% positive while the majority-Population 2 snails (Group B) are 41% positive. Total number of samples included in analysis was 503 including 496 genotyped-validation and genotyped-pooled-GWAS *B. sudanica* from this study, 5 inbred *B. sudanica* previously sequenced (27) and 2 outbred *B. choanomphala*. (B) Ordered Fisher’s exact test *p* values per variant of genotyped-validation samples within ancestry groups. *SudRes1* variants c6:3,490, c6:31,057, and c6844:72,816 are significant validated-variants (*p* < 1.3e-04, Bonferroni adjusted significance threshold shown by dotted line) within ancestry Group B (blue dots) and most other top-outliers (1.3e-04< *p* < 1.0e-03) are in *SudRes1* or *SudRes2*. (C) Ordered additive regression *p* values per variant of genotyped-validation samples, after accounting for ancestry. *SudRes2* variant sc94:2,166,296 is a significant validated-variant (*p* < 2.3e-04, Bonferroni adjusted significance threshold shown by dotted line) and most other top-outliers (1.3e-04< *p* < 1.0e-03) are in *SudRes1* or *SudRes2*. (D) Allele and genotype counts for the most significant validated-variants from *SudRes1* (c6:3,490) and *SudRes2* (sc94:2,166,296), which were both dominant markers, and the representative codominant marker variants from *SudRes1* (c582:65,596) and *SudRes2* (sc94:2,174,117), for *S. mansoni* infection positive (+) and negative (-) snails. Values are read counts for each allele in the pooled-GWAS (“GWAS”), and genotype counts in genotyped-validation snails (“Validation”). (E) Means and standard errors for each variable in the best-fitting multiple regression model. Ancestry is the proportion of Population 2 ancestry, and loci *SudRes1* and *SudRes2* are additive effects per each copy of the minor allele at the representative codominant markers c582:65,596 and sc94:2,174,117. \*\*\**p* < 0.001.

After accounting for ancestry, only variants within two genomic regions, henceforth referred to as *SudRes1* and *SudRes2*, showed significance (Bonferroni-corrected *p* < 0.05, designated henceforth as ‘validated-variants’) in a dominance (Fig. 2B) or an additive regression (Fig. 2C) model (Dataset S3). Notably, the p-values of the variants in *SudRes1* and *SudRes2* were amongst the lowest of the dual-variant outliers identified by the pooled-GWAS (Fig. 1). The three validated-variants for *SudRes1* (Fig. 2B) and validated-variant for *SudRes2* (Fig. 2C) acted as dominant markers (only two genotypes observed), so to assess genotype-phenotype associations in more depth we examined codominant proxy variants and used these as representative variants for each region (Fig. 2D). For the *SudRes1* representative variant (c582:65,596, Fig. 2D), allele T was protective in the pooled-GWAS and in the genotyped-validation snails. Combining data from the genotyped-validation and genotyped-pooled-GWAS snails, odds of *B. sudanica* infection with *S. mansoni* were 0.31 for genotype TT, 0.57 for genotype GT, and 1.37 for genotype GG, consistent with an additive effect. For the *SudRes2* representative variant (sc94:2,174,117, Fig. 2D), allele A was protective in the pooled-GWAS and in the genotyped-validation snails. Combining data from the genotyped-validation and genotyped-pooled-GWAS snails, odds of *S. mansoni* infection in *B. sudanica* were 0.36 for genotype AA, 0.74 for genotype TA, and 2.40 for genotype TT, also consistent with an additive effect.

The best-fitting multiple regression model (AIC = 451.11 and *p* < 0.001 for all variables) included: ancestry; *SudRes1* (representative variant c582:65,596, Fig. 2D); and *SudRes2* (representative variant sc94:2,174,117, Fig. 2D), with both genetic markers acting additively (Fig. 2E). The model predicts a ∼2-fold effect per allele at each genetic marker, and a ∼4-fold effect of ancestry. Thus, the predicted odds of infection for a snail with no Population 2 ancestry and major allele homozygous genotypes at *SudRes1* and *SudRes2* (4.46) is 62-fold higher (approximately 2^2^*2^2^*4) than the odds for a snail with 100% Population 2 ancestry and minor allele homozygous genotypes at both loci (0.07).

### SudRes1 is rich in paralogous genes encoding MEGF domains

*SudRes1* comprises 1.07 Mb of the *B. sudanica* genome and contains 23 protein coding genes across five contigs (c6844, c582, c5209, c6, c2) that are closely linked on chromosome 5 (Fig. 3A, Fig. S2 and Fig. S3). Notably, 9 of these 23 genes encode multiple epidermal growth factor (MEGF) domains (Fig. 3A, Dataset S4). Two of these MEGF proteins display a common single pass transmembrane domain (TMD) structure, with intracellular tyrosine-specific protein phosphatase (PTP) domains and extracellular MEGF and a galactose binding domain (GBD), forming a receptor-like PTP (RPTP) protein (Fig. 3B). Only 14 other MEGF/GBD-containing RPTP genes are annotated in the *B. sudanica* genome, 13 of which are clustered near *SudRes1* on chromosome 5 (Fig. 3A, Dataset S4).

**Fig. 3.**
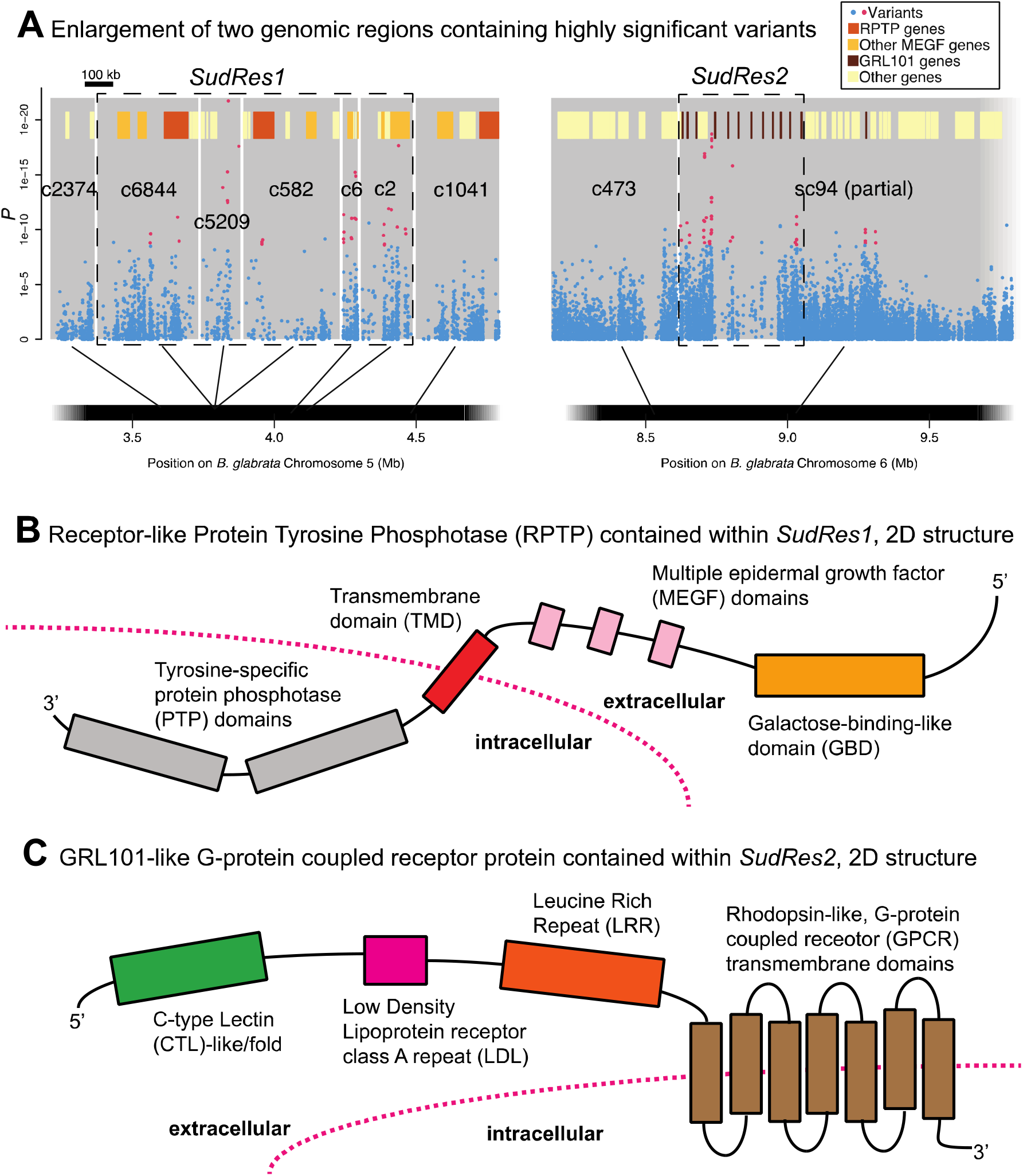
Characterization of *Biomphalaria sudanica SudRes1* and *SudRes2* genomic regions. (A) Pooled-GWAS *p* values in *SudRes1* and *SudRes2* regions (dashed boxes), which contain pooled-GWAS dual-variants (red) and other variants (blue), defined as in Figure 1. Contigs (grey rectangles) on *B. sudanica* chromosomes 5 and 6 are arranged horizontally based on contig orthology to 18 chromosomes of the *B. glabrata* genome (xgBioGlab47.1, NCBI RefSeq: GCF_947242115.1) and linkage map analysis (Dataset S2, Fig. S2-S4). Gene positions are shown (red/yellow/brown/orange boxes), highlighting three particularly prevalent classes of genes: in *SudRes1* the multiple epidermal growth factor (MEGF) and galactose-binding like domain (GBD) containing receptor-like tyrosine-specific protein phosphatase (RPTP), other protein coding genes containing MEGF domains, and; in *SudRes2* encoding a class of leucine-rich repeat containing G-protein couple receptors (GRL101) with C-type lectin and low-density lipoprotein extracellular domains (partial GRL101 genes included). (B) The predicted protein structure of receptor-like tyrosine-specific protein phosphatase (RPTP) coding gene BSUD.17727 (c6844) present in the *SudRes1* region of the *B. sudanica* genome and containing intronic validated-variants (Fig. S8A). A similar RPTP coding gene is contained within adjacent contig c582 within *SudRes1*, and contig c1041 neighboring *SudRes1* region (Fig. 3A). (C) A representative predicted protein structure of a GRL101-like G-protein coupled receptor coding gene, twelve of which were predicted through manual annotation within the *SudRes2* region of contig sc94 (1.82-2.26 Mb) in the *B. sudanica* genome.

Of the three validated-variants in *SudRes1* (Fig. 2B), one was contained within the intron of an MEGF/GBD-containing RPTP protein in contig c6844 (Fig. S8A), whilst the other two were in the intergenic region either side of another MEGF/GBD-containing protein (contig c6 ortholog 1, Dataset S4). Similarly, the two ‘top-outlier’ variants (defined as variants with 1.3e-04 < *p* < 1.0e-03 following validation, see Fig. 2B) in adjacent contigs c582 and c2 in *SudRes1* were contained within the intronic gene sequence of an MEGF/GBD-containing RPTP protein (c582 BSUD.15164, Dataset S4) and another MEGF/GBD-containing protein (c2 ortholog 2, Dataset S4).

Three of the five contigs in the *SudRes1* region, c6844, c5209 and c582, are homologous with each other and match the same unduplicated orthologous region on *B. glabrata* chromosome 5 and *B. pfeifferi* LG5 (Fig. S9 and S10). Aligned read coverage was also atypically low across all *SudRes1* contigs compared to the rest of the genome (Fig. S11).

### SudRes2 is characterized by a large family of GRL101-like GPCR genes

*SudRes2* comprises a 440 kb region between 1.82 and 2.26 Mb on contig sc94 of the *B. sudanica* chromosome 6 (Fig. 3A, Fig. S2 and Fig. S4). Following manual annotation of *SudRes2*, 14 protein coding genes were identified (Fig. S12 and Dataset S5). Ten of these encode mutually paralogous GRL101-like proteins, defined as G-protein coupled receptor (GPCR) transmembrane proteins with extracellular regions containing a leucine rich repeat (LRR) region, a low-density lipoprotein receptor class A repeat (LDL) and a C-type lectin-like (CTL) domain (Fig. 3D); an additional two GRL101-like genes in *SudRes2* are missing the CTL or LDL domain and may represent incomplete proteins (Dataset S5). Of the 437 GPCR genes in the *B. sudanica* genome (27), only six genes outside of *SudRes2* are annotated as possessing GRL101, CTL, and LDL domains (Dataset S5).

Both the validated-variant and top-outlier variant in *SudRes2* are contained in the non-coding regions of a gene which encodes a protein with a zinc finger RING-type (Zn-RING) domain and inhibitor of apoptosis (IAP) repeat region (i.e. Zn-RING-IAP) (Fig. S8B). This Zn-RING-IAP gene is within the cluster of GRL101 proteins in *SudRes2*. On the boundary of *SudRes2* is a baculoviral IAP repeat containing (BIRC) protein, many of which are contained in the genome regions neighboring *SudRes2*.

Prior to the manual annotation of these 14 genes, the *SudRes2* region was exceptional in that: 1) only four protein coding genes in 440 kb had been annotated in this region of the reference *B. sudanica* genome, much lower than the genome-wide density of one gene per 40 kb; 2) a low density of variants were present (Fig. 3A and Fig. S12), and; 3) a large drop in aligned pooled-GWAS read coverage across the central 240 kb of *SudRes2* was apparent (Fig. S13 and S14). To confirm the presence and validity of the manually annotated GRL101 genes, *B. sudanica* RNA transcript data was successfully aligned to 11 of the 12 predicted GRL101 CDS sequences in the *SudRes2* region (all except GRL101_3). Phylogenetic analysis of protein coding sequences also revealed that the *B. sudanica* syntenic (conserved gene order) orthologs identified in *B. glabrata* and *B. pfeifferi* (Dataset S5) were also the most closely related (Fig. S15).

## Discussion

### Identification of loci associated with schistosome resistance in a wild snail vector population

In this study, we identified and validated two previously uncharacterized genomic regions, *SudRes1* and *SudRes2*, in the African snail vector *B. sudanica* that are associated with resistance to *S. mansoni* infection, each contributing a similar effect size of a ∼2-fold change in *S. mansoni* infection odds ratio per allele. Both regions contain long segments with unusually low pooled-GWAS read coverage and contain few annotated genes in the reference genomes of *Biomphalaria* sp., suggesting possible structural variation or allelic divergence that preclude unambiguous alignment of reads and complicates assembly and annotation of these regions. It is crucial therefore to acknowledge that the validated-variants associated with schistosome resistance may not themselves be causal polymorphisms, instead they highlight that something significant is occurring in these regions that may remain elusive using the current *B. sudanica* genome assembly (27), potentially due to structure rearrangements or unaligned alleles. Manual annotation of both *SudRes* regions revealed that they are enriched with transmembrane protein coding genes with diverse extracellular regions composing of protein-protein interacting and carbohydrate binding domains, relevant to immune-related functions such as pathogen recognition (33, 34).

*SudRes1* is characterized by MEGF-domain containing genes, including receptor-like protein-tyrosine phosphatases (RPTPs), comprising extracellular MEGF, extracellular GBD, and intracellular tandem PTP domains. These potentially heavily glycosylated RPTPs may form stable dimers on the cell surface (35), with ligand binding triggering confirmational changes that expose or occlude catalytically active regions of the intracellular membrane-proximal PTPs, transducing signals across the cell membrane (36, 37). The presence of schistosome resistance-associated variants surrounding the *B. sudanica* RPTPs suggests that increased efficacy or upregulation of these proteins may counteract *S. mansoni*-induced phosphorylation, one of the parasite’s strategies to manipulate or evade the snail immune system and promote its survival (38). The *SudRes1* region was previously noted as showing exceptionally high nucleotide diversity in *B. sudanica* (27), which when coupled with the pooled-GWAS results suggest that pathogen-mediated balancing selection may act on these genes as previously hypothesized. Our findings here support the approach of using genome hyperdiversity as a proxy for identifying immune related genes in uncharacterized genomes.

*SudRes2* contains resistance-associated variants within a Zn-RING-IAP protein coding gene, which neighbors a cluster of 12 leucine rich repeat-containing G protein-coupled receptor (LGR) family proteins where pooled-GWAS variants are distributed throughout. The structure of the *SudRes2* LGR proteins is similar to GRL101, a LGR first described in the gastropod species *Lymnaea stagnalis* notable for its N-terminal extracellular LRRs and LDLs (UniProt accession P46023 (39)). Unique to the *B. sudanica* GRL101 genes characterized in *SudRes2*, however, is the N-terminal C-type lectin (CTL) fold/domain. CTL domain containing proteins are established components of both vertebrate and invertebrate innate immune systems as recognition and effector molecules, which show pathogen dependent expression patterns (40–42). Due to the architecture of the *SudRes2* GRL101 proteins, the CTL domain likely extends away from the cell membrane exposing the CTL binding region to cytoplasmic ligands (such as those derived from invading pathogens) that could then be presented to the GPCR membrane-spanning binding pocket, triggering G-protein activation. Homology and phylogenetic placement of the syntenic GRL101 proteins indicates a shared ancestry, and possible functional conservation, in GRL101 genes retained since the split of *B. glabrata* and African *Biomphalaria* species ∼5 Mya (26, 43), although currently available *Biomphalaria* sp. genome assemblies in this hyperdiverse region may impede inferences of expansion and contraction. To our knowledge, GRL101-like proteins have not been attributed to immunity in gastropods, but have been shown to play an important role in freshwater prawn innate immunity (44). Although GRL101-like genes were present elsewhere in the *B. sudanica* genome, the dense cluster of GRL101 genes in the *SudRes2* region is unique in that it is the only region <0.5 Mb with 12 GRL101 genes, with the caveat that GRL101 genes elsewhere in the *B. sudanica* genome may also not be annotated correctly.

Both *SudRes1* and *SudRes2* are evolutionarily dynamic. *SudRes1* shows evidence suggestive of either duplicated segment(s) relative to *B. glabrata* and *B. pfeifferi* or extremely divergent alleles that cannot be assembled, though close linkage across *SudRes1* was confirmed by a shared pattern of null genotypes in the linkage cross of F2 *B. sudanica* (Dataset S2). For *SudRes2*, the unusually low coverage indicates large deletions or high sequence divergence between individual snails, precluding confident read alignment and indicating a region of highly divergent haplotypes. Aligned coverage was also atypically low in whole-genome reads from four other *B. sudanica* inbred lines (27), though homologous reads in these data could be identified for many of the *SudRes2* GRL101 genes, and many of these genes were also present in the *B. glabrata* and *B. pfeifferi* genomes, showing some degree of intra- and inter-species conservation across this region.

### Evidence of a shifting snail population structure in Lake Victoria could lead to increased infections

A surprising result was the discovery of ancestry heterogeneity in our GWAS snails, whose parents had all been collected at the same time and place. More remarkable still, this ancestry signal is strongly correlated with schistosome resistance. Thus, many outliers in our pooled-GWAS could represent ancestry-informative markers with no physical linkage to resistance genes. Population ancestry estimates using only neutral linkage map markers were very similar to those using the full panel including GWAS outliers, supporting that the ancestry effect observed is real and not an artifact of using atypical variants implicated by the GWAS. Considering the importance of snail ancestry in schistosome compatibility here, potential causes behind the population structure were tested. First, no support for reproductively isolated cryptic *Biomphalaria* species in Lake Victoria causing the structure was found, since the ancestry estimates varied continuously between populations, and because no marker was fixed between ancestral populations. Second, since the estimated allele frequencies for the two ancestral populations are continuously distributed, and only few alleles are observed at a similar frequency, we do not expect that a single prolific snail had parented a disproportionate amount of the offspring used in the GWAS. Third, while *B. sudanica* and the deep-water taxon *B. choanomphala* are closely related, perhaps being ecophenotypes (32), sympatric, and distinct in parasite susceptibility (45), they do not represent the ancestry groups and cluster with *B. sudanica* having a majority Population 1 ancestry. Rather, Population 2 *B. sudanica* is distinguished by a set of alleles that do not appear to be common in either species.

The population structure of *B. sudanica* in Lake Victoria observed in our results is more consistent with historical isolation and reconnection of populations. Of the major lakes in the Albertine Rift Valley lake system (Victoria, Tanganyika, Malawi), Lake Victoria is a relatively young lake forming ∼0.4 Mya, and has gone through at least three major desiccations in the past 100,000 years (46–48). Shaped by such drought events, the cichlids of Lake Victoria have become a famous study system due to the astounding levels of explosive diversification that has occurred since the last desiccation event <15,000 years ago (49, 50). The population structure of *B. sudanica* observed suggests that indeed cryptic population structure is present, potentially caused by these historic events, yet the degree of admixing between populations in our study signifies ongoing outcrossing rather than clear speciation. The signatures of ongoing admixing may be influenced by hydrologic patterns of Lake Victoria. The collection site was ∼20 miles north of the Rusinga channel connecting the open lake and narrow Winam gulf. The Winam gulf is a unique lake environment given that it is somewhat separated from open lake water due to the prevailing currents limiting circulation of water (51, 52), is comparatively shallower, potentially exacerbating historic water level changes, and has more protected shores, providing different freshwater habitats than those present in the open lake. The hydrology of the Rusinga channel and therefore Winam gulf was most recently disrupted by the blocking of the Mbita passage in the early 1980’s, until its unblocking in 2017 (53), therefore occurring just prior to our snail collections in early 2018. The return of north-easternly flow of open-lake water into the Winam gulf through the Mbita passage has caused a shift in both bacterial and planktonic communities in the Winam gulf (54, 55), and may have allowed dispersal of *B. sudanica* populations on floating vegetation, such as water hyacinth between lake areas (56). Although we cannot establish the potentially different geographic origins of the *B. sudanica* representative of each population using currently available data, these snail population differences, and therefore vectoral competency differences, could explain why some locations around lakes are persistent hotspots of transmission while others are not (10). These findings underscore how pathogen resistance can vary substantially between closely related populations, and in this instance could suggest that schistosome transmission may be more persistent in lake regions where highly susceptible *Biomphalaria* populations, i.e. majority Population 1 ancestry, are present.

### Evolutionary dynamics of snail-schistosome interaction

This study complements extensive work on immune mechanisms in laboratory populations of *B. glabrata* (17, 20, 22, 57), facilitating comparisons between snail species. Notably, there was no overlap between our validated GWAS hits and loci linked to resistance in *B. glabrata*. The amplicon panel included at least two amplicons within or near each of these *a priori* candidates (27), and none of them showed a significant association with infection phenotype. *SudRes1* resides on chromosome 5, the site of a large resistance QTL in *B. glabrata* (22), though about 10 Mb away and thus unlikely to include the same gene(s). *SudRes2* resides on *B. sudanica* chromosome 6, which contains a density of schistosome-resistance *a priori* loci including *PTC1* (57), *tlr* (58), *sod1* (59), and the closest, *prx4* (60), at ∼1 Mb away is not likely to be responsible for association in our analysis. While allelic variation in orthologs of *B. glabrata* resistance loci are not associated with *S. mansoni* resistance in *B. sudanica*, reverse genetics approaches successfully applied in *B. glabrata* (61, 62) can be used in the future to functionally evaluate the roles of these genes. *Biomphalaria sudanica* may rely on entirely different genetic mechanisms for parasite resistance than *B. glabrata*. However, considering the extensive genotype-by-genotype interaction documented between *B. glabrata* and *S. mansoni* (63), mediated by hyperdiverse resistance loci suggestive of long-term balancing selection (17, 57), we propose a more nuanced scenario. Namely, that resistance alleles fluctuate dynamically in response to the genotypes of local parasites (including other trematodes), so the loci harboring large-effect, intermediate-frequency alleles will vary over time and space, even within a species. Other trematode species may be more prevalent and exert greater selection pressure on these snails, indirectly impacting resistance to *S. mansoni*. The striking 4-fold effect of ancestry on odds of infection also supports this dynamic view, as subpopulations different in resistance may be distributed unevenly across Lake Victoria. Furthermore, diversity at both *SudRes1* and *SudRes2* is high, potentially more so than we observe if nominally paralogous contigs at *SudRes1* are allelic and/or if low coverage at *SudRes2* is caused by extreme sequence divergence. Parasite-resistance regions *PTC1* (57) and *PTC2* (17) in *B. glabrata* are also highly polymorphic in *B. sudanica* (27), indicating a similar pattern of immune-relevant balancing selection consistent with long-term snail-parasite coevolution.

By revealing immune-relevant genetic variation in *B. sudanica*, the primary vector in African Great Lakes, this work represents an important step toward molecular-informed vector control to combat schistosomiasis in high-transmission global regions, including gene drive technologies (64). However, we demonstrate that genetic manipulation of snails for schistosomiasis control will require navigating the ever more complex genetic architecture of snail resistance, particularly when considering the non-overlap in findings from the laboratory model South American species *B. glabrata* and the diversity of snail vector species responsible for the majority of schistosome transmission in Sub-Saharan Africa. The impact of any resistance allele may vary due to genetic background and environmental factors. However, we expect that the large-effect loci identified here will play key roles in the continued elucidation of snail immunity as it pertains to human disease.

## Materials and Methods

Additional description of materials and methods is provided in SI Materials and Methods.

### Ethical considerations

This project was undertaken following approval from the relevant bodies, including Kenya Medical Research Institute (KEMRI) Scientific Review Unit (Approval # KEMRI/RES/7/3/1 and KEMRI/SERU/CGHR/035/3864), Kenya’s National Commission for Science, Technology, and Innovation (License # NACOSTI/P/15/9609/4270 and NACOSTI/P/22/14839), Kenya Wildlife Services (permit # 0004754 and # WRTI-0136–02-22), and National Environment, Management Authority (permit # NEMA/AGR/46/2014 – Registration # 0178 and NEMA/AGR/159/2022 – Registration # 201). Collections of *S. mansoni* from schoolchildren were approved by KEMRI’s Scientific and Ethics Review Unit (SERU), reference SERU No. 3540, and by the Institutional Review Board of the University of New Mexico (UNM), reference 18115. Informed consent was obtained from the parents of five children that were deemed to be positive for *S. mansoni* from fecal samples using egg microscopy, of which remaining samples were used for *S. mansoni* miracidial hatching. All five children were treated with praziquantel during a follow-up visit by a KEMRI physician.

## Data, Materials, and Software Availability

All sequence data has been uploaded onto the NCBI SRA under BioProject PRJNA1149315 with BioSample accessions SAMN43241892, SAMN43241893, SAMN43241894, SAMN43241895.

## Acknowledgments

This work was funded by National Institute of Health, National Institute of Allergy and Infectious Disease grants R01AI141862 and R37AI101438.

## References

1. WHO, Global report on neglected tropical diseases 2024 (2024).

2. S. I. Hay, et al., Global, regional, and national disability-adjusted life-years (DALYs) for 333 diseases and injuries and healthy life expectancy (HALE) for 195 countries and territories, 1990-2016: a systematic analysis for the Global Burden of Disease Study 2016. Lancet 390, 1260–1344 (2017).

3. WHO, “WHO guideline on control and elimination of human schistosomiasis” (2022).

4. WHO, “Ending the neglect to attain the Sustainable Development Goals – A road map for neglected tropical diseases 2021–2030” (2020).

5. N. C. Lo, et al., A call to strengthen the global strategy against schistosomiasis and soil-transmitted helminthiasis: the time is now. Lancet Infect. Dis. 17, e64–e69 (2017).

6. A. G. P. Ross, et al., A new global strategy for the elimination of schistosomiasis. Int. J. Infect. Dis. 54, 130–137 (2017).

7. S. H. Sokolow, et al., To Reduce the Global Burden of Human Schistosomiasis, Use ‘Old Fashioned’ Snail Control. Trends Parasitol. (2018).

8. N. Kittur, et al., Discovering, Defining, and Summarizing Persistent Hotspots in SCORE Studies. Am. J. Trop. Med. Hyg. 103, 24–29 (2020).

9. R. E. Wiegand, et al., A persistent hotspot of Schistosoma mansoni infection in a five-year randomized trial of praziquantel preventative chemotherapy strategies. J. Infect. Dis. 216, 1425–1433 (2017).

10. M. W. Mutuku, et al., A Search for Snail-Related Answers to Explain Differences in Response of Schistosoma mansoni to Praziquantel Treatment among Responding and Persistent Hotspot Villages along the Kenyan Shore of Lake Victoria. Am. J. Trop. Med. Hyg. 101, 65–77 (2019).

11. H. F. Tavalire, M. S. Blouin, M. L. Steinauer, Genotypic variation in host response to infection affects parasite reproductive rate. Int. J. Parasitol. 46, 123–131 (2016).

12. C. H. King, D. Bertsch, Historical Perspective: Snail Control to Prevent Schistosomiasis. PLoS Negl. Trop. Dis. 9, e0003657 (2015).

13. S. H. Sokolow, et al., Global Assessment of Schistosomiasis Control Over the Past Century Shows Targeting the Snail Intermediate Host Works Best. PLoS Negl. Trop. Dis. 10, e0004794 (2016).

14. W. E. Secor, Water-based interventions for schistosomiasis control. Pathog. Glob. Health 108, 246–254 (2014).

15. D. J. Civitello, et al., Transmission potential of human schistosomes can be driven by resource competition among snail intermediate hosts. Proc. Natl. Acad. Sci. 119, e2116512119 (2022).

16. C. M. Adema, et al., Will all scientists working on snails and the diseases they transmit please stand up? PLoS Negl. Trop. Dis. 6, e1835 (2012).

17. J. A. Tennessen, et al., Clusters of polymorphic transmembrane genes control resistance to schistosomes in snail vectors. Elife 9, e59395 (2020).

18. J. Buchthal, S. W. Evans, J. Lunshof, S. R. Telford, K. M. Esvelt, Mice Against Ticks: an experimental community-guided effort to prevent tick-borne disease by altering the shared environment. Philos. Trans. R. Soc. B Biol. Sci. 374, 20180105 (2019).

19. J. R. Powell, Modifying mosquitoes to suppress disease transmission: Is the long wait over? Genetics 221, iyac072 (2022).

20. L. Lu, L. Bu, S.-M. Zhang, S. K. Buddenborg, E. S. Loker, An Overview of Transcriptional Responses of Schistosome-Susceptible (M line) or -Resistant (BS-90) Biomphalaria glabrata Exposed or Not to Schistosoma mansoni Infection. Front. Immunol. 12, 805882 (2022).

21. C. M. Adema, et al., Whole genome analysis of a schistosomiasis-transmitting freshwater snail. Nat. Commun. 8, 15451 (2017).

22. L. Bu, et al., Compatibility between snails and schistosomes: insights from new genetic resources, comparative genomics, and genetic mapping. Commun. Biol. 5, 940 (2022).

23. J. A. Tennessen, et al., Genome-Wide Scan and Test of Candidate Genes in the Snail Biomphalaria glabrata Reveal New Locus Influencing Resistance to Schistosoma mansoni. PLoS Negl. Trop. Dis. 9, 1–19 (2015).

24. M. Knight, et al., The identification of markers segregating with resistance to Schistosoma mansoni infection in the snail Biomphalaria glabrata. Proc. Natl. Acad. Sci. 96, 1510–1515 (1999).

25. P. J. Hotez, A. Kamath, Neglected Tropical Diseases in Sub-Saharan Africa: Review of Their Prevalence, Distribution, and Disease Burden. PLoS Negl. Trop. Dis. 3, 1–10 (2009).

26. L. Bu, et al., A genome sequence for Biomphalaria pfeifferi, the major vector snail for the human-infecting parasite Schistosoma mansoni. PLoS Negl. Trop. Dis. 17, e0011208 (2023).

27. T. Pennance, J. Calvelo, et al, The genome and transcriptome of the snail Biomphalaria sudanica s.l.: Immune gene diversification and highly polymorphic genomic regions in an important African vector of Schistosomas mansonis. BMC Genomics 25, 192 (2024).

28. H. Bastide, et al., A Genome-Wide, Fine-Scale Map of Natural Pigmentation Variation in Drosophila melanogaster. PLoS Genet. 9, e1003534 (2013).

29. T. Pennance, et al., A rapid diagnostic PCR assay for the detection of Schistosoma mansoni in their snail vectors. J. Parasitol. in press.

30. N. R. Campbell, S. A. Harmon, S. R. Narum, Genotyping-in-Thousands by sequencing (GT-seq): A cost effective SNP genotyping method based on custom amplicon sequencing. Mol. Ecol. Resour. 15, 855–867 (2015).

31. D. H. Alexander, J. Novembre, K. Lange, Fast model-based estimation of ancestry in unrelated individuals. Genome Res. 19, 1655–1664 (2009).

32. P. S. Andrus, J. R. Stothard, N. B. Kabatereine, C. M. Wade, Comparing shell size and shape with canonical variate analysis of sympatric Biomphalaria species within Lake Albert and Lake Victoria, Uganda. Zool. J. Linn. Soc. 139, 713–722 (2023).

33. N. Dinguirard, et al., Proteomic Analysis of Biomphalaria glabrata Hemocytes During in vitro Encapsulation of Schistosoma mansoni Sporocysts. Front. Immunol. 9, 2773 (2018).

34. X.-J. Wu, et al., Proteomic analysis of Biomphalaria glabrata plasma proteins with binding affinity to those expressed by early developing larval Schistosoma mansoni. PLoS Pathog. 13, 1–30 (2017).

35. G. Jiang, J. den Hertog, T. Hunter, Receptor-Like Protein Tyrosine Phosphatase α Homodimerizes on the Cell Surface. Mol. Cell. Biol. 20, 5917–5929 (2000).

36. J. N. Andersen, et al., Structural and Evolutionary Relationships among Protein Tyrosine Phosphatase Domains. Mol. Cell. Biol. 21, 7117–7136 (2001).

37. N. K. Tonks, Protein tyrosine phosphatases: from genes, to function, to disease. Nat. Rev. Mol. Cell Biol. 7, 833–846 (2006).

38. A. J. Walker, D. Rollinson, Specific tyrosine phosphorylation induced in Schistosoma mansoni miracidia by haemolymph from schistosome susceptible, but not resistant, Biomphalaria glabrata. Parasitology 135, 337–345 (2008).

39. C. P. Tensen, et al., A G protein-coupled receptor with low density lipoprotein-binding motifs suggests a role for lipoproteins in G-linked signal transduction. Proc. Natl. Acad. Sci. 91, 4816–4820 (1994).

40. B. Pees, et al., Effector and regulator: Diverse functions of C. elegans C-type lectin-like domain proteins. PLoS Pathog. 17, e1009454 (2021).

41. H. Schulenburg, M. P. Hoeppner, J. Weiner, E. Bornberg-Bauer, Specificity of the innate immune system and diversity of C-type lectin domain (CTLD) proteins in the nematode Caenorhabditis elegans. Immunobiology 213, 237–250 (2008).

42. I. M. Dambuza, G. D. Brown, C-type lectins in immunity: recent developments. Curr. Opin. Immunol. 32, 21–27 (2015).

43. R. J. DeJong, et al., Phylogeography of Biomphalaria glabrata and B. pfeifferi, important intermediate hosts of Schistosoma mansoni in the New and Old World tropics. Mol. Ecol. 12, 3041–3056 (2003).

44. X. Cao, et al., Leucine-rich repeat-containing G-protein-coupled receptor 2 (LGR2) can regulate PO activity and AMP genes expression in Macrobrachium nipponense. Mol. Immunol. 126, 14–24 (2020).

45. M. W. Mutuku, et al., Comparative vectorial competence of Biomphalaria sudanica and Biomphalaria choanomphala, snail hosts of Schistosoma mansoni, from transmission hotspots In Lake Victoria, Western Kenya. J. Parasitol. 107, 349–357 (2021).

46. T. C. Johnson, K. Kelts, E. Odada, The holocene history of Lake Victoria. Ambio 2–11 (2000).

47. J. C. Stager, T. C. Johnson, The late Pleistocene desiccation of Lake Victoria and the origin of its endemic biota. Hydrobiologia 596, 5–16 (2008).

48. A. S. Cohen, M. J. Soreghan, C. A. Scholz, Estimating the age of formation of lakes: An example from Lake Tanganyika, East African Rift system. Geology 21, 511–514 (1993).

49. F. Ronco, et al., Drivers and dynamics of a massive adaptive radiation in cichlid fishes. Nature 589, 76–81 (2020).

50. C. Müller, F. N. Moser, D. Frei, O. Seehausen, Constraints to speciation despite divergence in an old haplochromine cichlid lineage. Evolution (N. Y). 77, 823–835 (2023).

51. P. Okely, J. rg Imberger, J. P. Antenucci, Processes affecting horizontal mixing and dispersion in Winam Gulf, Lake Victoria. Limnol. Oceanogr. 55, 1865–1880 (2010).

52. C. Nyamweya, et al., Simulation of Lake Victoria Circulation Patterns Using the Regional Ocean Modeling System (ROMS). PLoS One 11, e0151272 (2016).

53. B. M. Simiyu, H. S. Amukhuma, L. Sitoki, W. Okello, R. Kurmayer, Interannual variability of water quality conditions in the Nyanza Gulf of Lake Victoria, Kenya. J. Great Lakes Res. 48, 97–109 (2022).

54. B. M. Simiyu, R. Kurmayer, Response of planktonic diatoms to eutrophication in Nyanza Gulf of Lake Victoria, Kenya. Limnologica 93, 125958 (2022).

55. K. M. Brown, et al., Bacterial community and cyanotoxin gene distribution of the Winam Gulf, Lake Victoria, Kenya. Environ. Microbiol. Rep. 16, e13297 (2024).

56. M. L. Plummer, Impact of Invasive Water Hyacinth (Eichhornia crassipes) on Snail Hosts of Schistosomiasis in Lake Victoria, East Africa. Ecohealth 2, 81–86 (2005).

57. J. A. Tennessen, et al., Hyperdiverse Gene Cluster in Snail Host Conveys Resistance to Human Schistosome Parasites. PLoS Genet. 11, 1–21 (2015).

58. E. A. Pila, M. Tarrabain, A. L. Kabore, P. C. Hanington, A novel Toll-like receptor (TLR) influences compatibility between the gastropod Biomphalaria glabrata, and the digenean trematode Schistosoma mansoni. PLoS Pathog. 12 (2016).

59. C. P. Goodall, R. C. Bender, J. K. Brooks, C. J. Bayne, Biomphalaria glabrata cytosolic copper/zinc superoxide dismutase (SOD1) gene: Association of SOD1 alleles with resistance/susceptibility to Schistosoma mansoni. Mol. Biochem. Parasitol. 147, 207–210 (2006).

60. M. K. Larson, R. C. Bender, C. J. Bayne, Resistance of Biomphalaria glabrata 13-16-R1 snails to Schistosoma mansoni PR1 is a function of haemocyte abundance and constitutive levels of specific transcripts in haemocytes. Int. J. Parasitol. 44, 343–353 (2014).

61. E. A. Pila, et al., Endogenous growth factor stimulation of hemocyte proliferation induces resistance to Schistosoma mansoni challenge in the snail host. Proc. Natl. Acad. Sci. 113, 5305–5310 (2016).

62. P. C. Hanington, et al., Role for a somatically diversified lectin in resistance of an invertebrate to parasite infection. Proc. Natl. Acad. Sci. 107, 21087–21092 (2010).

63. J. P. Webster, M. E. J. Woolhouse, Cost of resistance: relationship between reduced fertility and increased resistance in a snail-schistosome host-parasite system. Proc. R. Soc. London B Biol. Sci. 266, 391–396 (1999).

64. T. Maier, et al., Gene drives for schistosomiasis transmission control. PLoS Negl. Trop. Dis. 13, 1–21 (2019).

