## Supplemental Information and Figures for "Immune targets for schistosomiasis control identified by a genome-wide association study of East African snail vectors"

**Other supporting materials for this manuscript include the following:**

Datasets S1 to S5

**SI Materials and Methods**

**Snail phenotyping.** In total, 329 *Biomphalaria sudanica* snails were collected from Anyanga Beach Kanyibok, Kenya (Latitude: -00.08958°, Longitude: 34.08592°) on March 27^th^ 2018. These snails were screened the following day for any patent echinostome infections, of which eight were found and discarded, leaving 321 snails that formed the parental generation. Parent snails were bred in plastic aquaria in an outdoor rearing facility at Kenya Medical Research Institute (KEMRI) in Kisumu. Snails housed in aquaria were allowed to breed and lay eggs for 7-14 days, then the adults moved to a new aquarium so that the eggs can hatch, and juveniles develop. This process was repeated every 7-14 days for approximately two months to reach the goal of 1400 offspring juvenile snails with a shell diameter of 4-6 mm to use in our infection cohort, which was reached in May 2018.

*Schistosoma mansoni* eggs were obtained from five local school children aged 6-12 years old that attended the regional primary school, Kanyibok Primary School, Usenge, Kenya (Latitude: -00.08552°, Longitude: 34.07778°), on May 30^th^ 2018, following informed consent from parents. Infected children were given praziquantel (40 mg/kg body weight) treatment during a follow-up visit by a KEMRI physician. Individual fecal samples were screened for *S. mansoni* eggs using the Kato-Katz method and then five samples with high egg intensity were selected for use in snail challenges that were performed over the course of the next two days, storing fecal samples at 4° C overnight. The eggs hatch upon exposure to freshwater and light and the resulting free-swimming miracidia were used for snail exposures.

In total, 1400 offspring juvenile snails (4-6 mm) were each exposed to eight freshly-hatched miracidia in the wells of a 24-well tissue culture plates each filled with ~2ml of bottled water overnight. Exposed snails were maintained in large plastic aquaria (~20L) in an open-air snail facility under ambient light and fed lettuce. Each snail was checked for infection via cercarial shedding every three to five days from 5- to 9-weeks post exposure. Shedding snails were considered positive and persevered in 100% ethanol. Any snails that did not release *S. mansoni* cercariae by 9-weeks post-exposure were later subjected to a PCR diagnostic to ensure they were truly negative (uninfected) as opposed to harboring a prepatent/latent infections (1). Only snails that were cercariae and PCR negative were considered negative and used for GWAS.

**Pooled genome-wide association study.** Genomic DNA (gDNA) from each snail was extracted from headfoot tissue following a modified CTAB protocol for freshwater snails (2). For the positive and negative pools, 10 ng of gDNA from each snail was included in the appropriate pool after quantification by two Qubit DS DNA assays (Invitrogen, CA, USA). Two Nextera libraries were created per pool as technical replicates to help ensure robustness and mitigate potential biases, and each was subjected to paired end 150 bp sequencing across two lanes of the Illumina NovaSeq 6000 S4 flow cell, with the positive and negative snail libraries on one lane and the technical replicate libraries of each on the other.

All custom *perl* scripts used in the following bioinformatic analysis are available in the project repository: github.com/jacobtennessen/PoolG. To process the resulting DNA sequence read data from each pool, Nextera Illumina Adapters were cut in *cutadapt* v3.1 (3) and reads timed in *trimmomatic* using options: LEADING:20 TRAILING:20 SLIDINGWINDOW:5:20 MINLEN:50) (4). *Schistosoma mansoni* reads were then removed from the DNA libraries by aligning data to the *S. mansoni* v9 genome (GCA_000237925.5 (5) using *bwa mem* (6, 7) and removing reads that mapped with CIGAR value >70 using a custom *perl* script (*FilterMappedBamCigar.pl,* options: -m 70) whilst removing secondary and supplementary reads in *samtools* using options: -f 1 -F 2304 -bS (8) to retain only reads that are *B. sudanica*. FASTQ files of the filtered *B. sudanica* reads were generated using *BEDtools* function *bamtofastq* (9) then aligned to the *B. sudanica* genome (10) using *bwa mem* (6, 7). PCR duplicate reads were marked, and removed, along with unmapped, secondary and supplementary alignments, and only paired reads were kept (*samtools* (8) options: view -f 1 -F 3332 -bS). Due to large file size, some files had to be partitioned for improved processing, these were merged using *samtools merge*, before each bam file representing each library was indexed; *samtools index*.

The allele depth was used to calculate allele counts and therefore frequencies to analyze genotype-phenotype associations between our positive and negative pools of snails. First a Variant Call Format (VCF) file was generated using the *bcftools* (11) *mpileup* (options: -A -a AD) and *call* (options -m -Ov) functions. Subsequently, a custom *perl* script that extracted information from the allele depth region (*AlleleCountsFromGwasVcf.pl*, options: -d 20 -q 0.05 -l 0,1 -s 100 -i), was used to make a table of counts of each allele for each sample in the VCF. The options correspond to filtering the variant data by using a minimum allele count of 20, and removing variants if any of the four quality metrics (RPB, MQB, MQSB, and BQB) contained in the VCF were <0.05, or if allele depth was <100. Indels were removed. This resulted in a table of counts of each allele for each sample in the VCF. A Chi-squared test was used to test for enrichment of outlier *p* values for variants between technical replicates. Technical replicates were then combined evenly in downstream analysis due to observed similarity.

For each variant in the *B. sudanica* genome that passed the filtering process, we calculated Fishers exact test *p* values from 2x2 contingency tables representing allele count by phenotype (positive or negative), and the odds ratios for each variant, in R v4.3 (12). This resulted in a table denoting one row per variant with columns providing the allele count in positive and negative pools, odds ratio, Fisher’s and Chi-squared *p* values. This was then summarized using a custom *perl* script (*SummarizeGWAS_p2.pl*).

*B. sudanica* contigs/scaffolds were ordered (Fig. 1) based on orthology to *B. glabrata* iM scaffolds and linkage groups (13), following our previous approach (10). Namely, we used BLASTP of all *B. glabrata* proteins against all *B. sudanica* proteins, and vice versa, to identify reciprocal best hits. *B. sudanica* contigs/scaffolds for which a majority of ortholog-matched genes occurred on the same *B. glabrata* scaffold were assigned a position based on the location of those orthologs.

**Validation of variants by amplicon genotyping.** To validate the pooled-GWAS outliers, a multiplex amplicon panel was designed using the Genotyping-in-Thousands by sequencing (GT-Seq) method (14). The panel contains 470 amplicons in total, including markers for 234 dual-variants i.e. genomic locations with two or more proximate variants (<50 kb) that were strong outliers (Fishers exact test *p* < 2.5e-9), and 12 singleton-variants (with *p* < 1e-13 from the pooled-GWAS), and 22 markers for *a priori* gene candidates based on variants located near candidate immune genes of *B. glabrata* or exceptionally diverse regions of *B. sudanica* (10). We applied the panel to the 220 independent validation snails (122 positive, 98 negative) and 276 of the pooled-GWAS snails (138 positive, 138 negative), referred to as genotyped-validation and genotyped-pooled-GWAS snails respectively, to confirm genotypes of snail individuals and validate the differential allele frequencies observed in the pooled-GWAS sequencing data.

In addition to the pooled-GWAS variants and *a priori* gene markers were an additional 201 ‘neutral’ markers (i.e. with no expected phenotypic association) included in the amplicon panel to facilitate the inclusions of large contigs on a linkage map (see **Linkage mapping**, below). Also, a *Schistosoma* 16S amplicon to confirm parasite presence was included in the panel for future applications since a PCR diagnostic (1) was used to determine infected snails in this study.

In addition to the GWAS positive and negative *B. sudanica* genotyped by the amplicon panel, *t*wo outbred *B. choanomphala* snails collected in Lake Victoria using offshore deep-water dredging in 2019 from Ndiara Beach (Latitude:-00.090925°, Longitude:34.085067°) were also included for purposes of investigating population structure and utility of the developed panel on this taxon.

Amplicon genotyping was conducted by GTseek (Twin Falls, Idaho), which reports read counts of each allele per sample (14). Amplicon genotypes were inferred from read counts as follows: <20x coverage considered missing, if each allele has >10% the coverage of the other allele the genotype is designated heterozygote, otherwise the genotype is designated homozygote. We assessed population structure in genotyped-validation snails and genotyped-pooled-GWAS snails using ADMIXTURE (15) and principal component analysis (PCA) in R. We also performed PCA including genotypes from the two outbred *B. choanomphala* samples, and the genotypes of five previously sequenced *B. sudanica* inbred line snail genomes (10). Given a signal of population structure, we accounted for ancestry in our validation analysis using two complementary statistical approaches. First, we fit logistic regression model for each marker in the set of genotyped-validation samples, assuming an additive effect on phenotype, and including ancestry as a quantitative cofactor (proportion of ancestry from second population). We designated significance using a Bonferroni threshold of 0.05/N, where N is the number of candidate amplicons (i.e. excluding linkage map amplicons) with minor allele frequency >5% and ≥150 non-missing genotypes in genotyped-validation samples. Second, for a dominance model we converted genotype markers to two sets of binary variables, alternately designating heterozygotes as 0 or 1, divided genotyped-validation samples into two groups based on ancestry (group A and B), and performed Fisher’s exact tests between genotypes and phenotypes within each ancestry group. We designated a significance threshold as before, this time accounting for two tests per marker per ancestry group and requiring non-missing data at 80 samples.

To find the best-fitting model relating genotype to phenotype from the amplicon panel data, we combined both genotyped-validation and genotyped-pooled-GWAS snail data. We tested all combinations of logistic multiple regression models encompassing ancestry and all successfully validated loci, considering all combinations of additivity and dominance at the genetic markers. We chose the model with the lowest AIC, and for which all individual variables have *p* < 0.05.

**Linkage mapping.** To generate crossed snails for linkage mapping, two juvenile (<4 mm) *B. sudanica*, one from each line 163 and KEMRI, which demonstrate opposite patterns of susceptibility to *S. mansoni* (UNMKenya line) (16), were reared in a ~1 liter breeding tank. Once egg laying and hatched juveniles were present in the tank, the two parent snails and eight randomly selected juveniles had gDNA extracted using the Qiagen Blood & Tissue kit (Qiagen, MD, USA). Outcrossing was confirmed using a developed PCR and restriction enzyme based diagnostic marker (17). Remaining F1 juveniles were moved to a new tank and then reared until eggs and F2 juveniles were present. These F2 juveniles were reared until >3 mm size. gDNA of the Bs163 and BsKEMRI parents and 87 F2 *B. sudanica* snails were genotyped using the amplicon panel as described above.

Genotype data from parents and F2s were used to create genomic linkage maps with OneMap v.3.0.0 (18), using minimum logarithm of the odds (LOD) of 4.5 from the rf_2pts option to unite markers into linkage groups. We included all amplicon markers that segregated in the F2s, whether they were included as GWAS variants or linkage map markers. Amplicon sequences were then matched to the *B. glabrata* genome assembly (xgBioGlab47.1, NCBI RefSeq GCF_947242115.1) using BLAST, facilitating synteny comparison between our *B. sudanica* map and the *B. glabrata* genome.

**Characterization of validated genomic regions**

Regions of the *B. sudanica* genome that were significantly associated with *S. mansoni* resistance and then validated were characterized using the *B. sudanica* genome annotation (10) to determine the function of protein coding genes, and annotate any potentially missing genes from the current *B. sudanica* annotation. Synteny of regions to the *B. glabrata* genome assembly (xgBioGlab47.1, NCBI RefSeq GCF_947242115.1) and *B. pfeifferi* genome assembly UNM_Bpfe_1.0 (GenBank GCA_030265305.1) were assessed using D-GENIES (19). tBLASTn searches against the *B. sudanica* genome were conducted using proteins in the syntenic regions of the *B. glabrata* genome. Alignments of *B. glabrata* and *B. pfeifferi* homologs/orthologs (see Dataset S4 and S5) to *B. sudanica* sequences, were used where possible to determine complete gene sequences in the case that the reference protein sequences were suspected to be truncated. Open reading frames (ORFs) were assessed in the *B. sudanica* validated genomic regions using the in-built tool in Geneious v2022.0.2 (Biomatters Ltd.) and predicted protein sequences generated that were characterized based on the protein families and functional domains predicted using InterProScan (20) and DeepTMHMM (21). Neighboring ORFs in the same direction were merged (internal stop codons in ORFs, predicted to be in intronic regions, were removed) to construct complete coding sequences of suspected proteins. Available PacBio and Illumina RNA-seq transcript data for *B. sudanica* (NCBI accessions: SRX22544968- SRX22544970) were aligned using *bwa mem* (6) to predicted mRNA sequences of these proteins.

BLASTn searches of predicted mRNA sequences generated following manual annotation were performed against the *B. sudanica* genome to determine any other homologous genes in the genome that may have not been annotated. For the 23,598 genes assigned to have open reading frames in the *B. sudanica* genome (10), these were searched against using key descriptors of genes of interest (i.e. PTPRA and GR101). In addition, InterPro accessions attributed to each gene in the annotated *B. sudanica* genome were searched against InterPro gene families and functional domains of interest. Specifically, to determine homologous genes to GRL101-like G-protein coupled receptor (GRL101) proteins containing a leucine rich repeat (LRR) region, a low-density lipoprotein receptor class A repeat (LDL) and a C-type lectin-like (CTL) domain, we filtered gene data by those matching InterPro accessions: IPR000276; IPR017452; IPR001304; IPR016187; IPR016186; IPR032675; IPR001611; IPR036055; IPR002172. To determine homologous genes to receptor-like protein-tyrosine-specific phosphatases (RPTP) containing multiple epidermal growth factor (MEGF) and galactose binding domain(s) (GBD), we filtered gene data by those matching InterPro accessions: IPR000242; IPR02902; IPR016130; IPR000387; IPR003595; IPR008979; IPR009030; IPR000742; IPR002049.

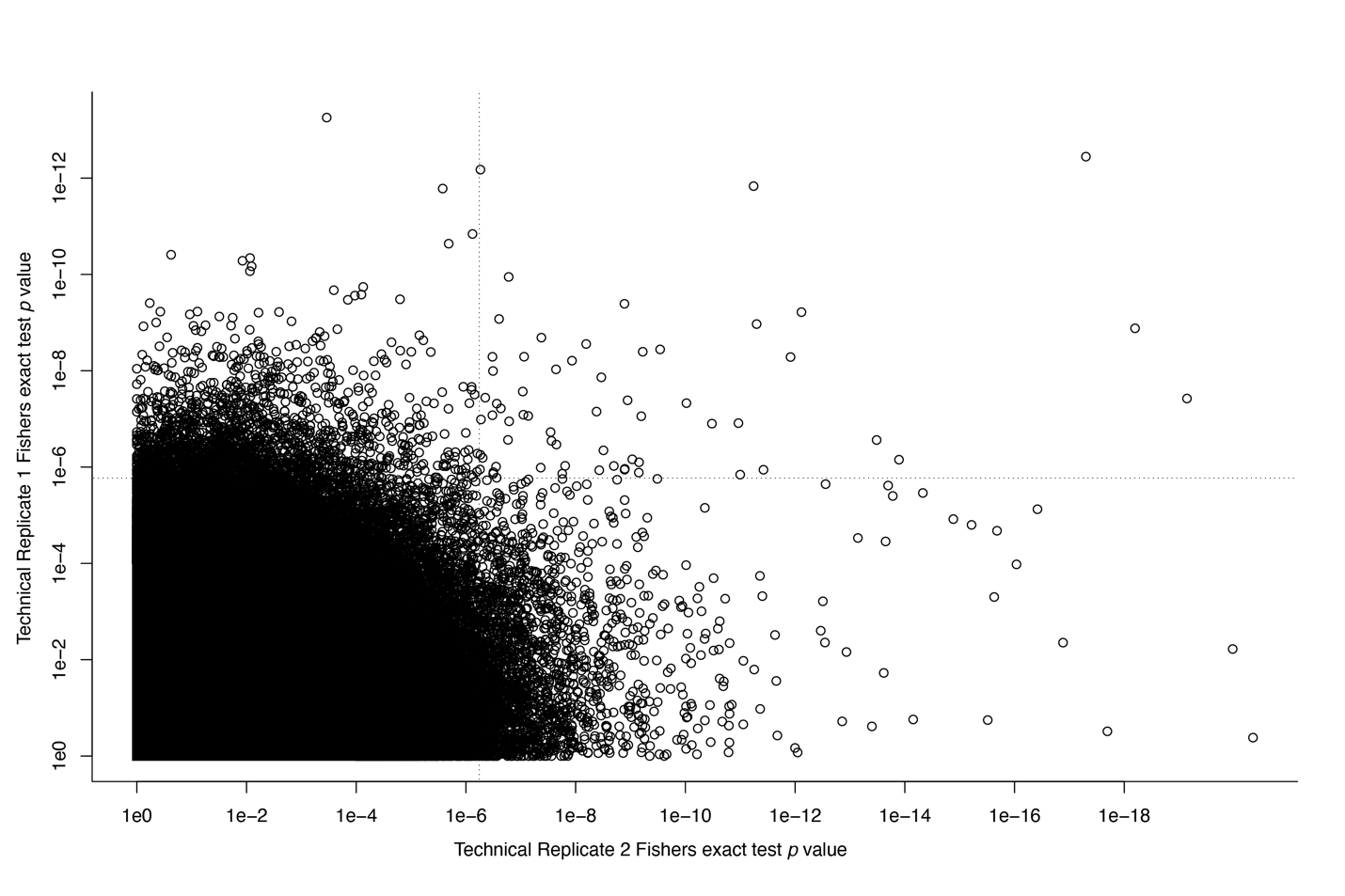

**Fig. S1.** Plot of Fisher exact test *p* values of variants across both pooled-GWAS technical replicates, showing a weak but significantly positive correlation (*R^2^* = 0.015). The dotted lines in the top right quadrant mark the region which the expected number of dots is 1 if there were no correlation.

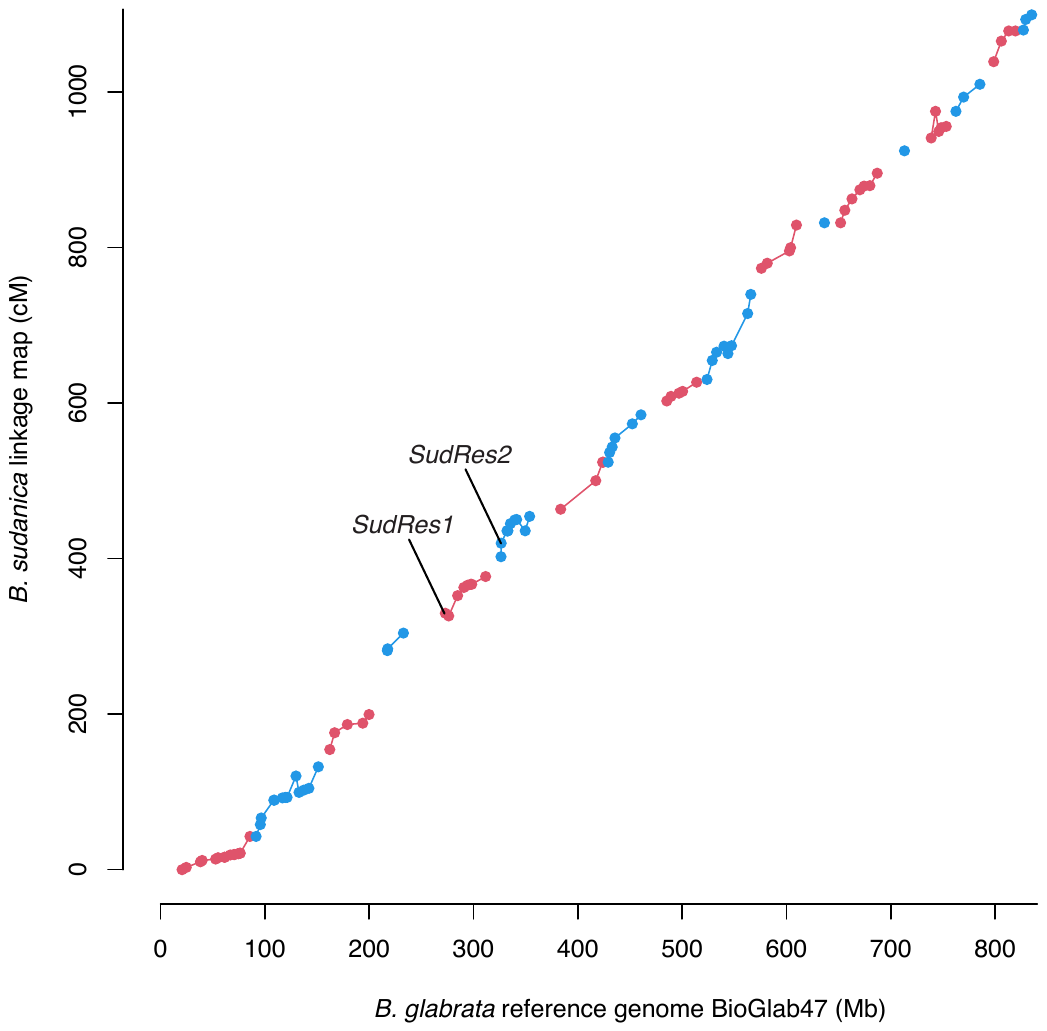

**Fig. S2.** *Biomphalaria sudanica* linkage map generated from 87 *B. sudanica* F2s generated from two parents from *B. sudanica* inbred lines 163 and KEMRI (16), aligned to *B. glabrata* reference genome xgBioGlab47.1 (Accession GCF_947242115.1). Chromosomes/linkage groups are alternately colored blue and red. The locations of the two validated GWAS loci associated with schistosome resistance in *B. sudanica* in the current study, *SudRes1* and *SudRes2*, are indicated.

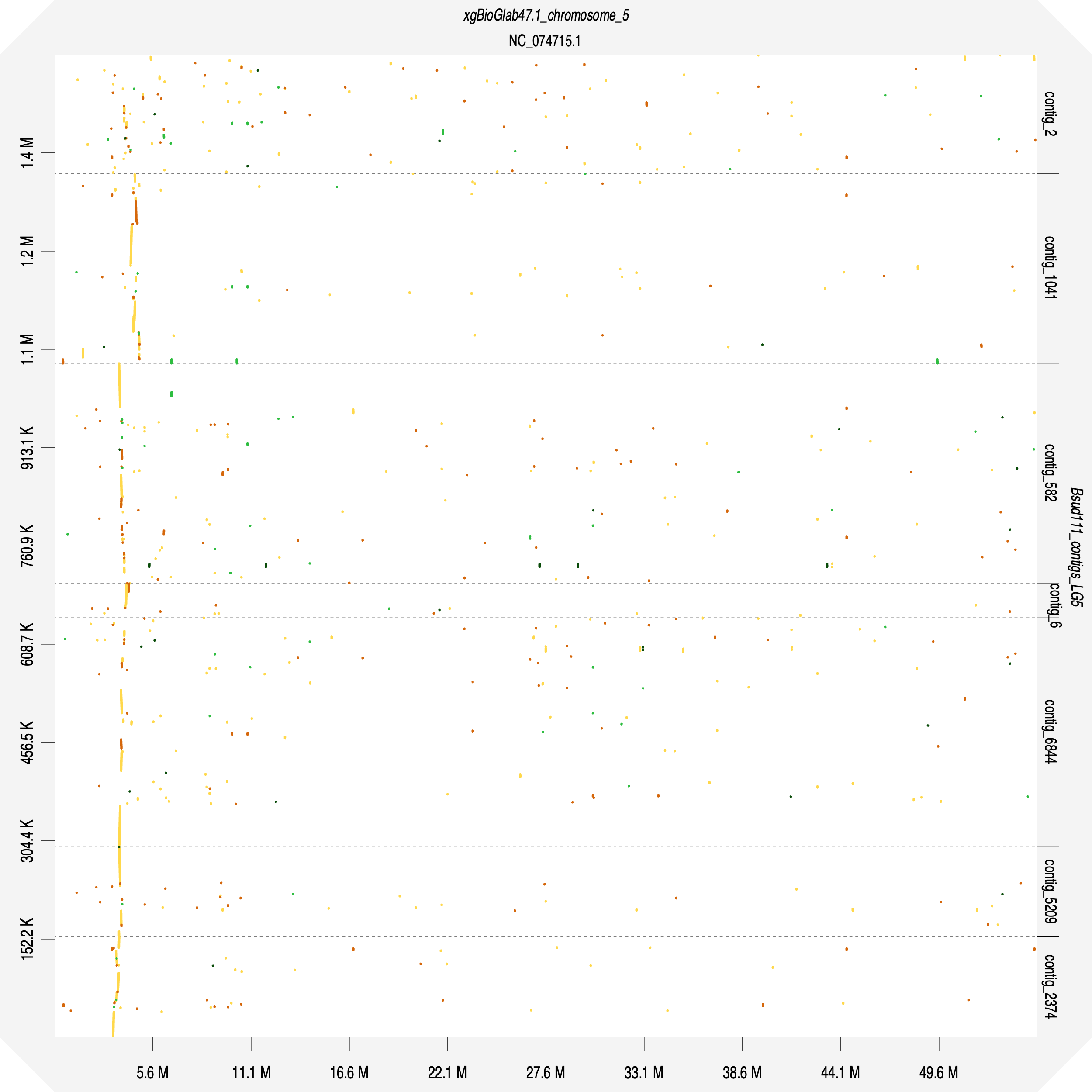

**Fig. S3.** Dot plot constructed using D-GENIES (19) comparing *Biomphalaria sudanica* genome (10) *SudRes1* region and its neighboring contigs against the *B. glabrata* genome (xgBioGlab47.1, Accession GCF_947242115.1) complete chromosome 5 sequence. Line color represents D-GENIES identity: yellow = 0 to <0.25; orange = 0.25 to <0.50; light green = 0.5 to <0.75; dark green = 0.75 to 1.

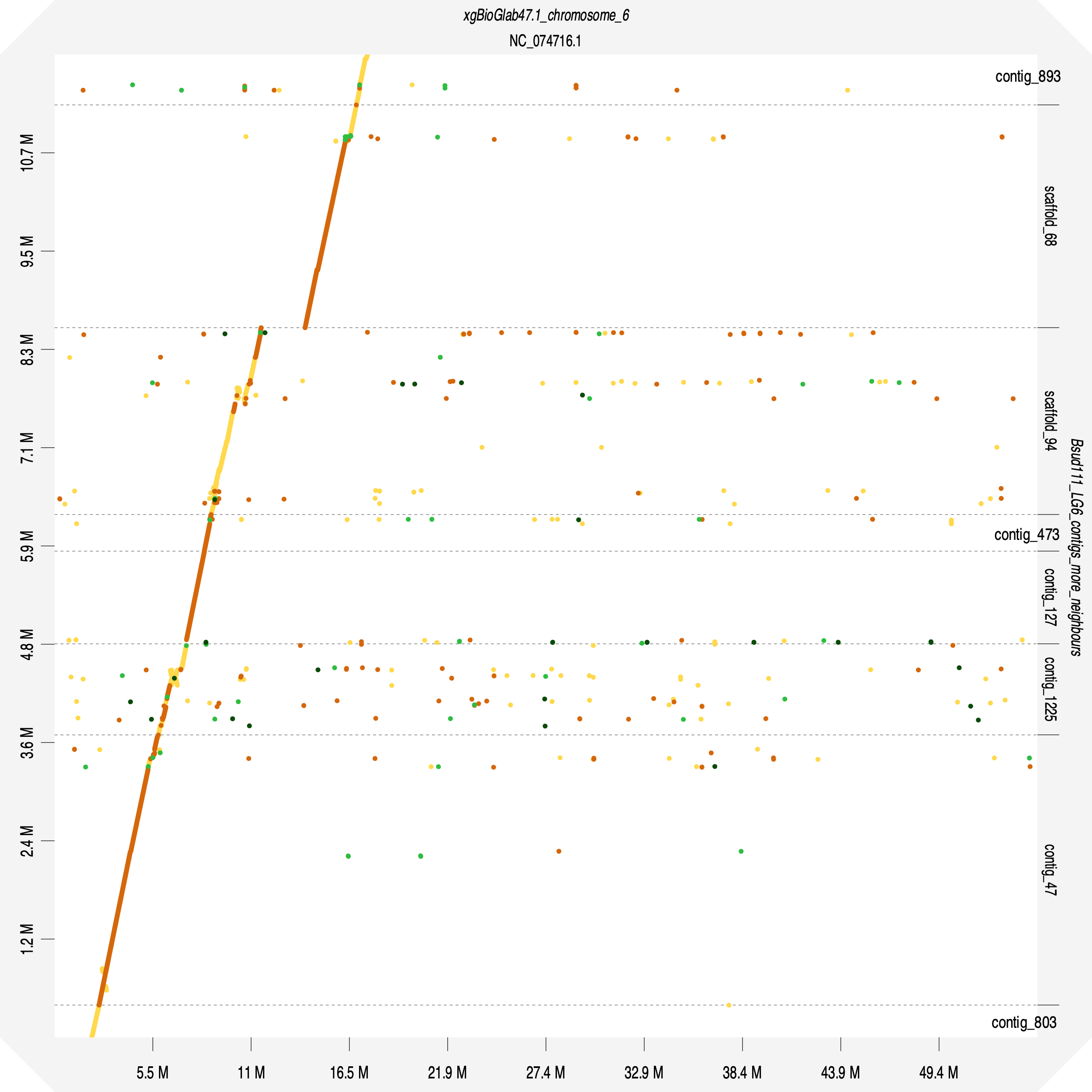

**Fig. S4.** Dot plot constructed using D-GENIES (19) comparing *Biomphalaria sudanica* genome (10) *SudRes2* region (contig sc94: 1.82 – 2.26 Mb) and its neighboring regions/contigs against the *Biomphalaria glabrata* genome xgBioGlab47.1 (Accession GCF_947242115.1) complete chromosome 6 sequence. Line color represents D-GENIES identity: yellow = 0 to <0.25; orange = 0.25 to <0.50; light green = 0.5 to <0.75; dark green = 0.75 to 1.

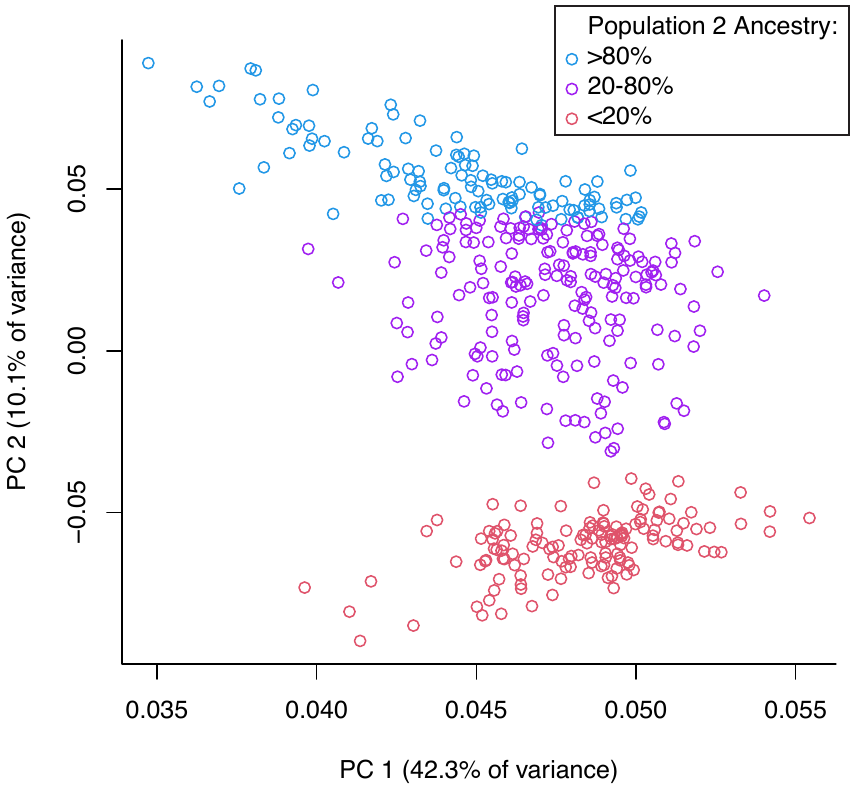

**Fig. S5.** Principal Component Analysis of amplicon genotypes recapitulates ancestry inference from ADMIXURE (15). We included all 457 loci with <50% missing data in redone and validation samples, and all 451 individuals with <10 missing genotypes at these loci (missing data imputed as heterozygous). Samples are colored based on Population 2 Ancestry results of ADMIXTURE (15) analysis.
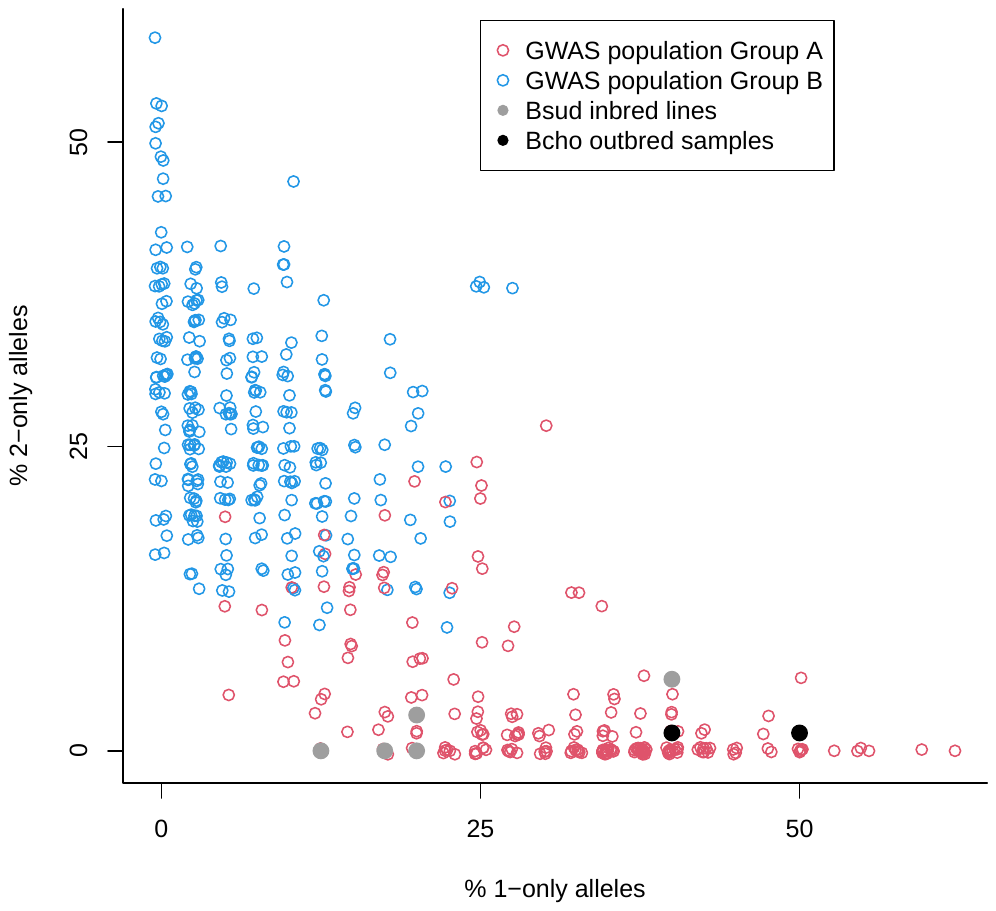

**Fig. S6.** Comparison of ADMIXTURE (15) results for GWAS *B. sudanica* with two outbred *B. choanomphala* (Bcho, black circles) and five previously sequenced (10) *B. sudanica* inbred lines (Bsud, grey circles). As ADMIXTURE estimates no fixed differences between ancestral populations 1 and 2, we examined alleles which are absent in one population (≤0.001%) and common in the other population (>20%), comprising 20 “1-only alleles” (absent in Population 2) and 34 “2-only alleles” (absent in Population 1). In *B. choanomphala*, 14 (70%) of the 1-only alleles were observed while only 2 (6%) of the 2-only alleles were observed. Similarly, in *B. sudanica* inbred lines, 15 (75%) of the 1-only alleles were observed while only 4 (12%) of the 2-only alleles were observed. In contrast, GWAS Group B snails (majority Population 2 ancestry, blue) have mostly 2-only alleles. Thus, both *B. sudanica* and *B. choanomphala* are a much closer match to Group A (majority Population 1 ancestry, red) at these diagnostic markers, and the population structure in the GWAS snails is not reflective of this interspecies difference.

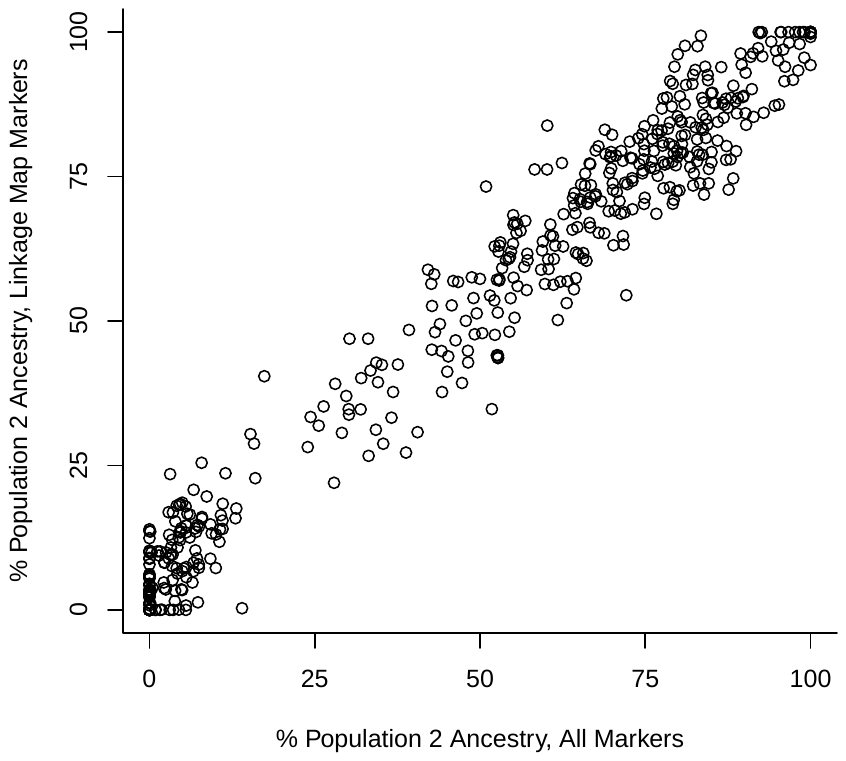

**Fig. S7.** Comparison of ADMIXTURE (15) results using all amplicon panel markers (x-axis) or only linkage map markers (y-axis) excluding GWAS candidates: dual- and singleton-variants and *a priori* loci (Dataset S3).

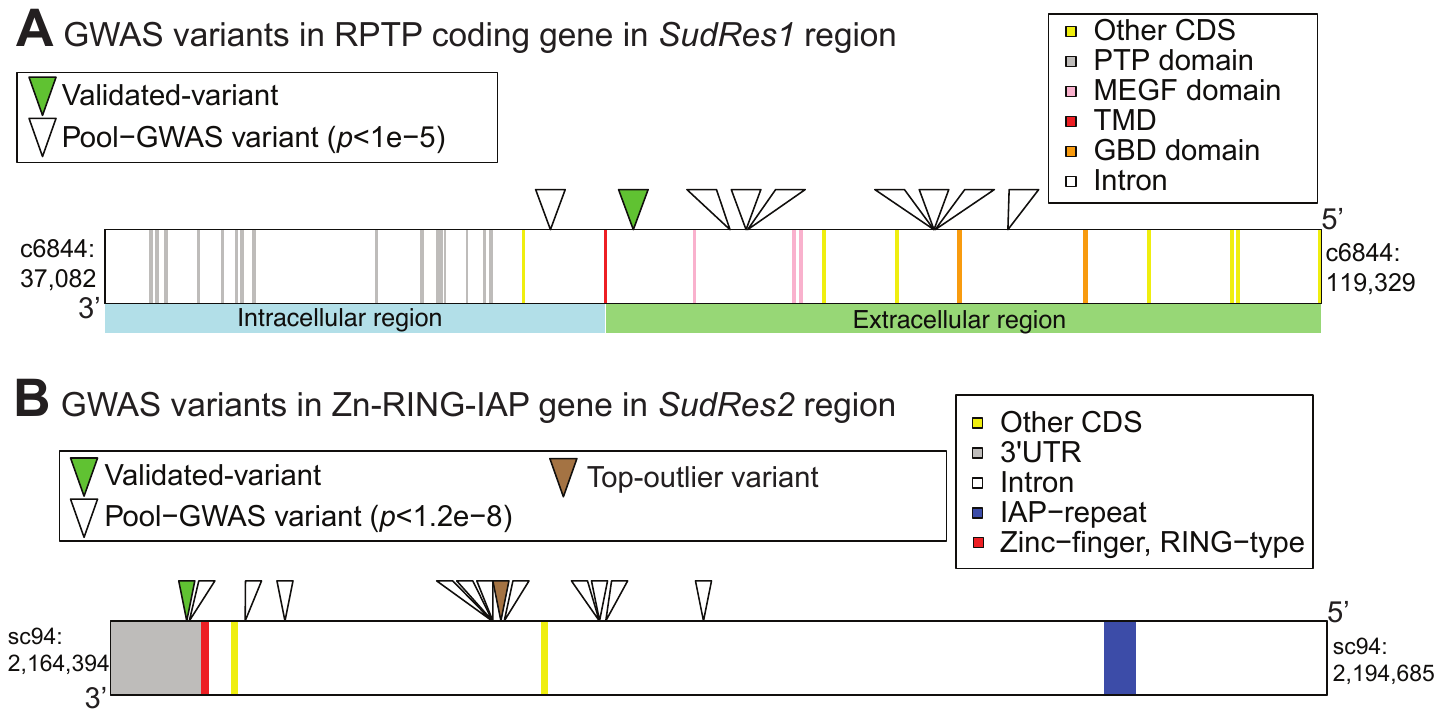

**Fig S8.** Position of validated-variants in *SudRes1* and *SudRes2* relative to full gene sequences, including introns, coding sequence (CDS) and CDS functional domains. (A) Plot showing nucleotide positions of CDS and CDS functional domains of a receptor-like tyrosine-specific protein phosphatase (RPTP) coding gene (BSUD.17727, Dataset S4) present in the *SudRes1* region of the *B. sudanica* genome. CDS of single pass transmembrane domain (TMD), multiple epidermal growth factor (MEGF) and galactose-binding like (GBD) domains are shown. Also shown are the nucleotide position of the validated-variant in c6844 (see Fig. 2B) and surrounding pooled-GWAS variants with *p* < 1e-5, all in introns. (B) Plot showing nucleotide positions of CDS and CDS functional domains within Zinc-finger-RING-type, inhibitor of apoptosis repeat (Zn-RING-IAP) containing gene (BSUD.25704, Dataset S5). Also shown are the nucleotide position of the validated-variant and top-outlier variant (see additive regression model Fig. 2C) and surrounding pooled-GWAS variants with *p* < 1.2e-8 (28 other variants with *p* < 1e-5 not shown for display purposes), all in introns and 3’UTR regions.

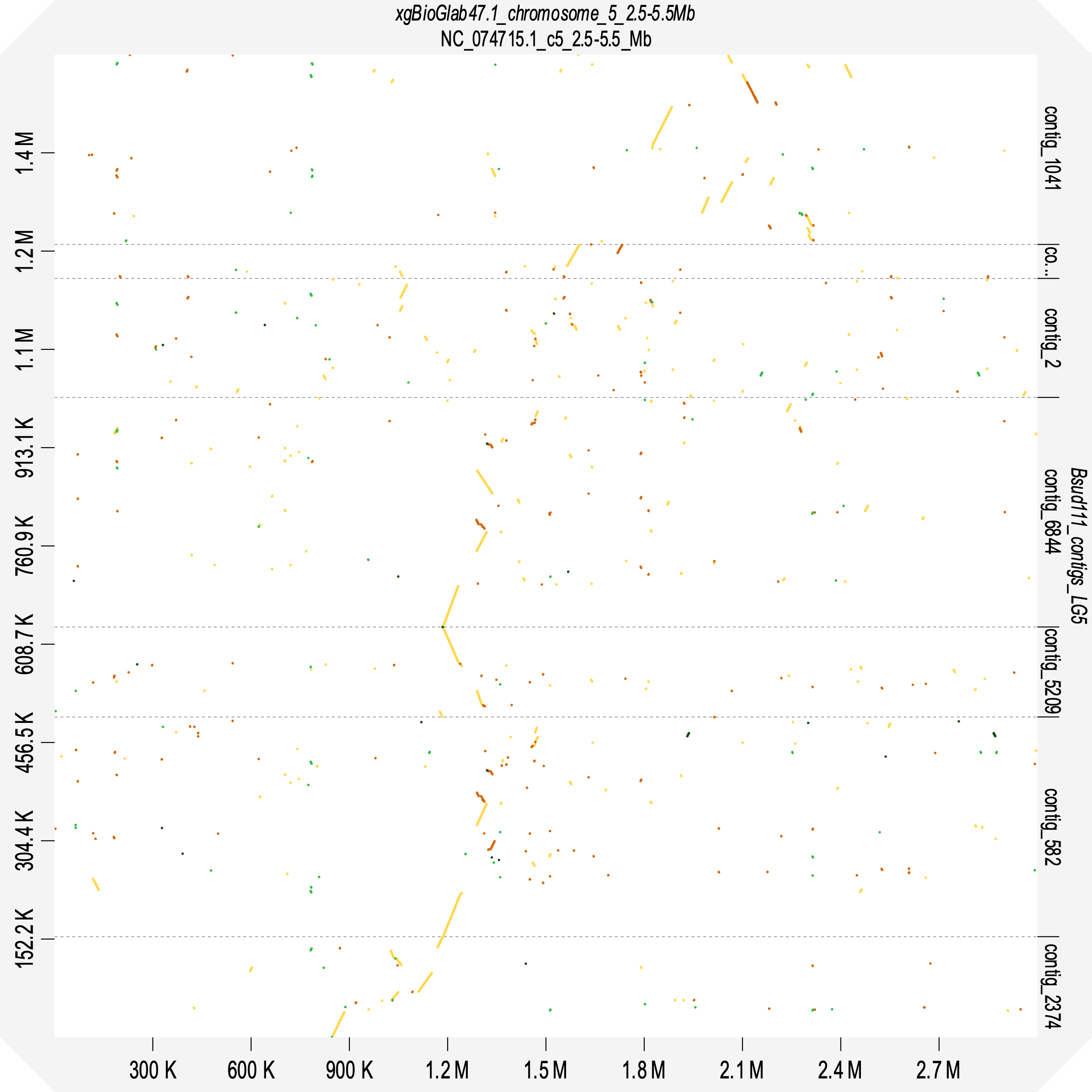

**Fig. S9.** Dot plot constructed using D-GENIES (19) comparing synteny of *Biomphalaria sudanica* genome (10) *SudRes1* region (contigs c6884, c582, c5209, c6, c2) and neighboring contigs (c1041 and c2374) against the *Biomphalaria glabrata* genome (xgBioGlab47.1, Accession GCF_947242115.1) 2.5 – 5.5 Mb region of chromosome 5. Line color represents D-GENIES identity: yellow = 0 to <0.25; orange = 0.25 to <0.50; light green = 0.5 to <0.75; dark green = 0.75 to 1.

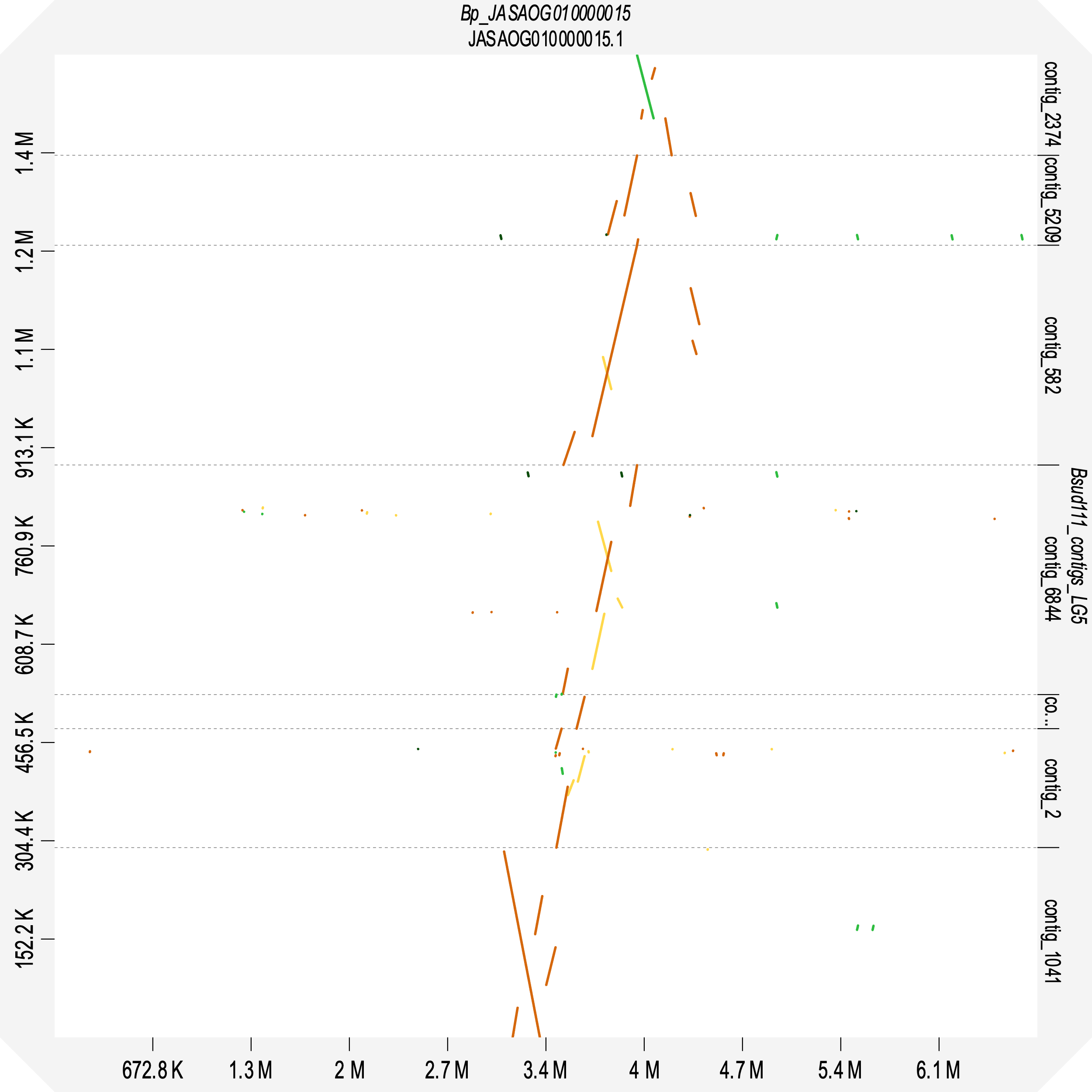

**Fig. S10.** Dot plot constructed using D-GENIES (19) showing synteny of *Biomphalaria sudanica* genome (10) *SudRes1* region (contigs c6884, c582, c5209, c6, c2) and neighboring contigs (c1041 and c2374) against the *Biomphalaria pfeifferi* genome (UNM_Bpfe_1.0, GenBank GCA_030265305.1) orthologous contig JASAOG010000015. Line color represents D-GENIES identity: yellow = 0 to <0.25; orange = 0.25 to <0.50; light green = 0.5 to <0.75; dark green = 0.75 to 1.

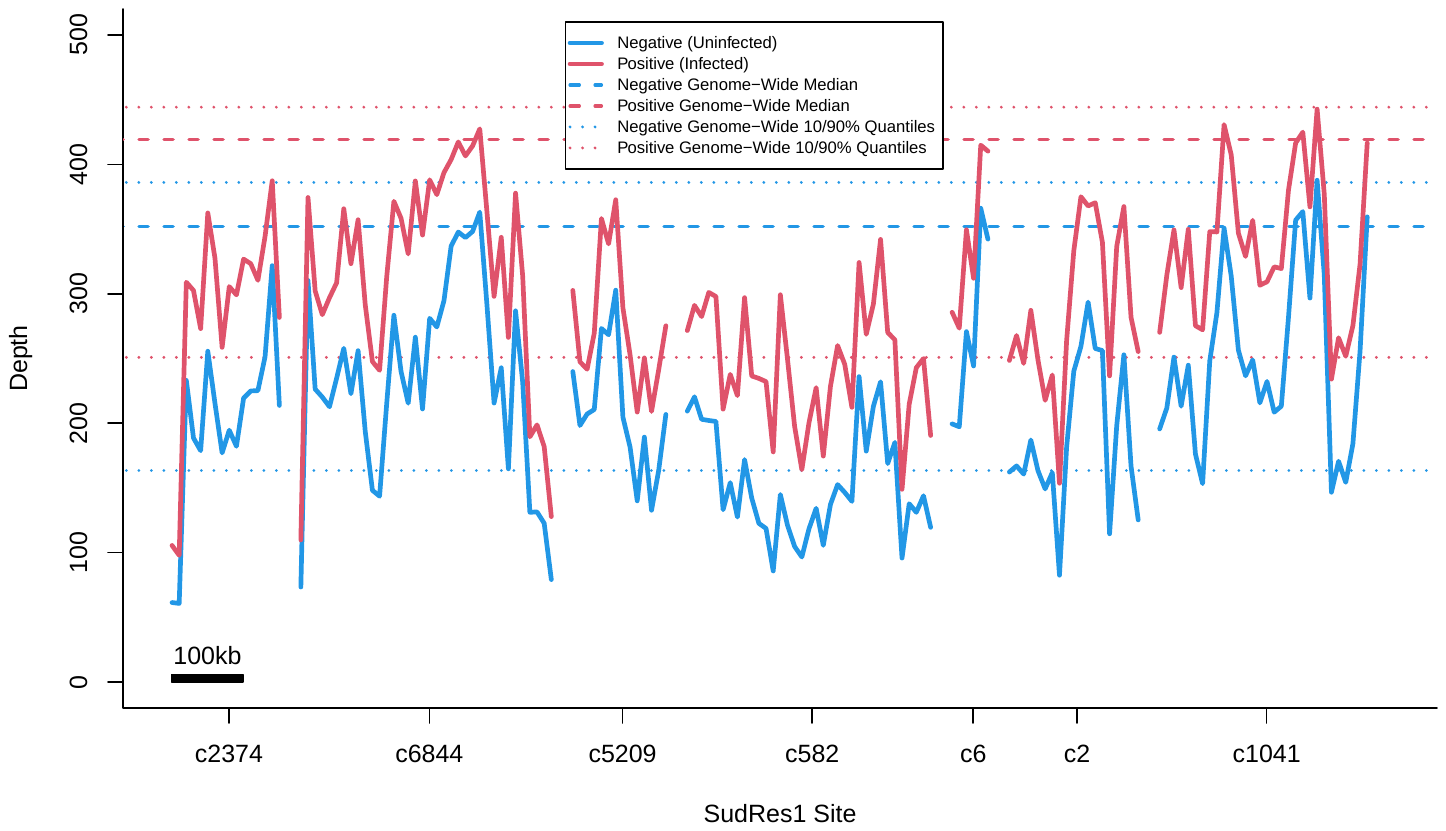

**Fig. S11.** Aligned read coverage, shown as mean read depth, in 10 kb windows of Pooled-GWAS data for the Negative (blue) and Positive (red) pools (both technical replicates combined) in the *SudRes1* genomic region (c6844, c5209, c582, c6, c2) and neighboring contigs. Across all *SudRes1* contigs, coverage is relatively low, often less than the 10% quantile for the genome (dotted lines).

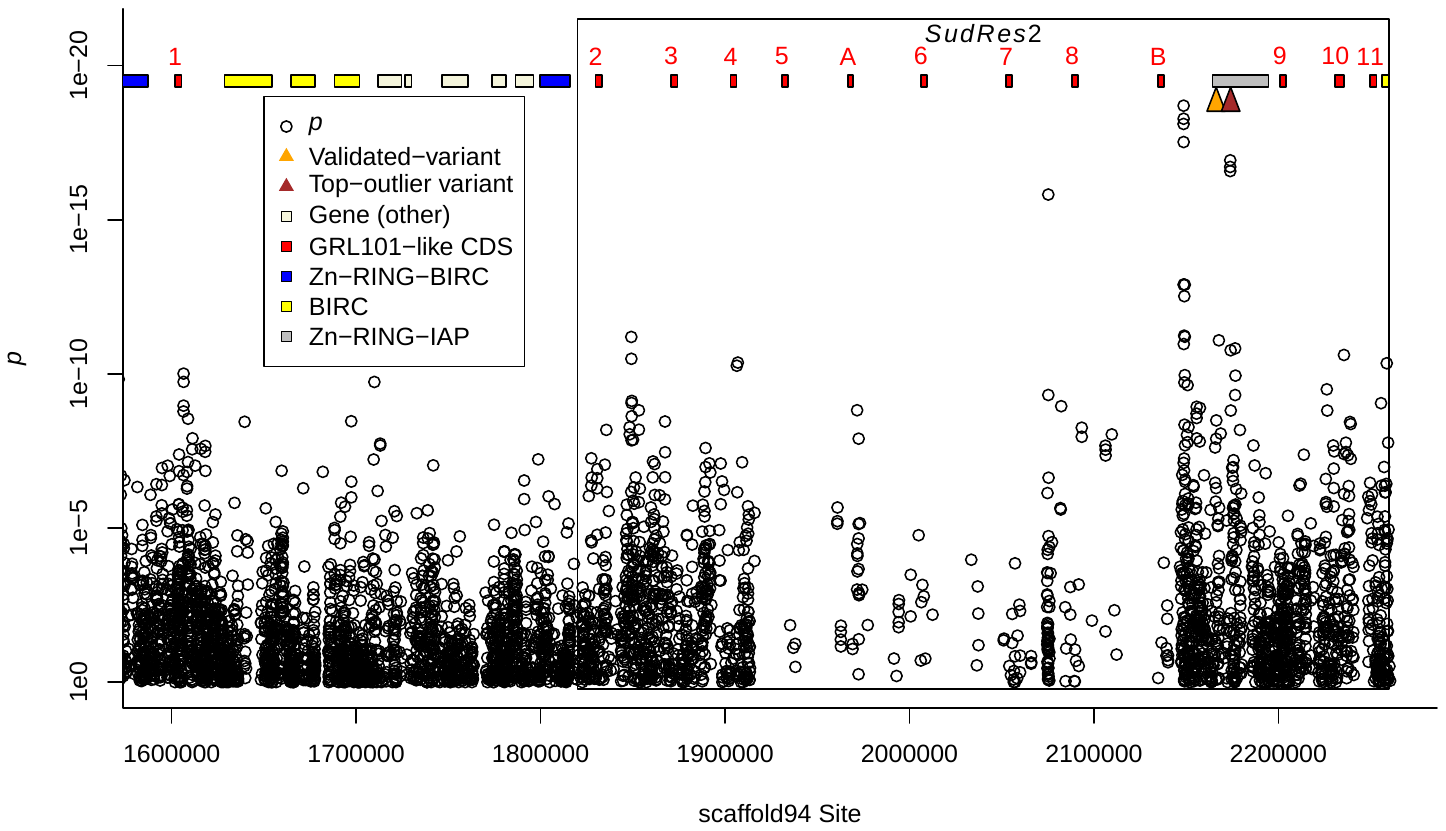

**Fig. S12.** Enlargement of contig sc94 (1.6 Mb to end, displayed in forward direction) on *Biomphalaria sudanica c*hromosome 6, showing Fisher’s exact test *p* values for all pooled-GWAS variants. The *SudRes2* region (represented by the black box surrounding sc94:1.82-2.26 Mb), contains 23 pooled-GWAS dual-variants with Fisher’s test *p* ≤ 5e-11, as well as the validated-variant (sc94:2,166,296, see Fig. 2C and 2D) and top-outlier variant (sc94:2,174,117, see Fig. 2C and 2D) associated with *B. sudanica* resistance to *Schistosoma mansoni*. Positions of the 13 (12 in *SudRes2*) GRL101-like G protein coupled receptor coding sequences (GRL101-like CDS) in sc94 are shown, as well as positions of genes containing combinations of zinc finger RING-type (Zn-RING), inhibitor of apoptosis (IAP) and baculoviral IAP repeat containing (BIRC) domains.

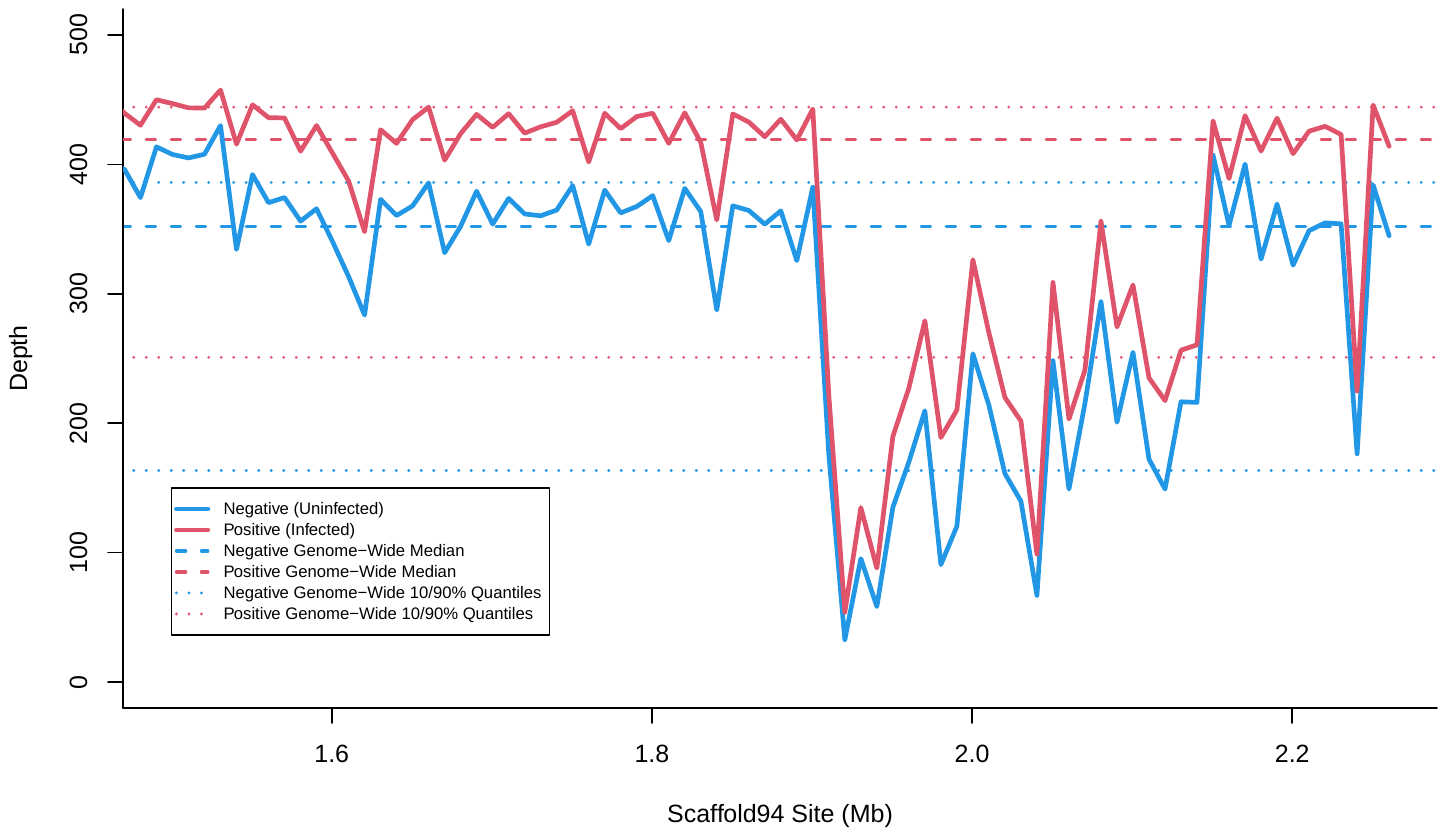

**Fig. S13**. Aligned read coverage, shown as mean read depth, in 10 kb windows of Pooled-GWAS data for the Negative and Positive pools (both technical replicates combined). The *SudRes2* gene region where a cluster of GRL101-like GPCR proteins were manually annotated lies between 1.6 Mb to the end of the contig sc94, corresponding to the region where read depth drops.

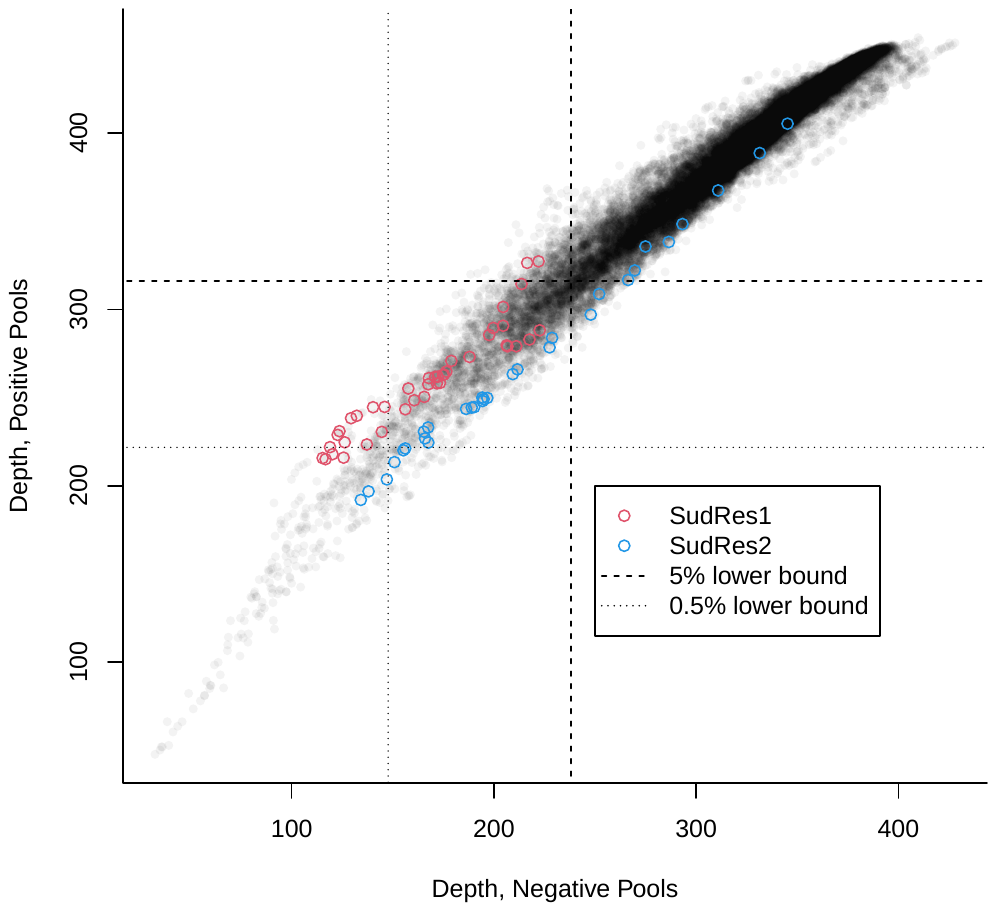

**Fig. S14.** Genome-wide pooled-GWAS mean depths in sliding 100 kb windows (step size = 10 kb) for the negative and positive pools (both technical replicates combined). Both *SudRes1* and *SudRes2* are notable for showing unusually low depth as well as slightly skewed ratios of positive/negative depth (unusually high for *SudRes1* and unusually low for *SudRes2*). Low coverage may indicate these regions are particularly variable and dynamic, with large sequence difference from the reference genome and/or poor mappability among highly similar adjacent paralogs.

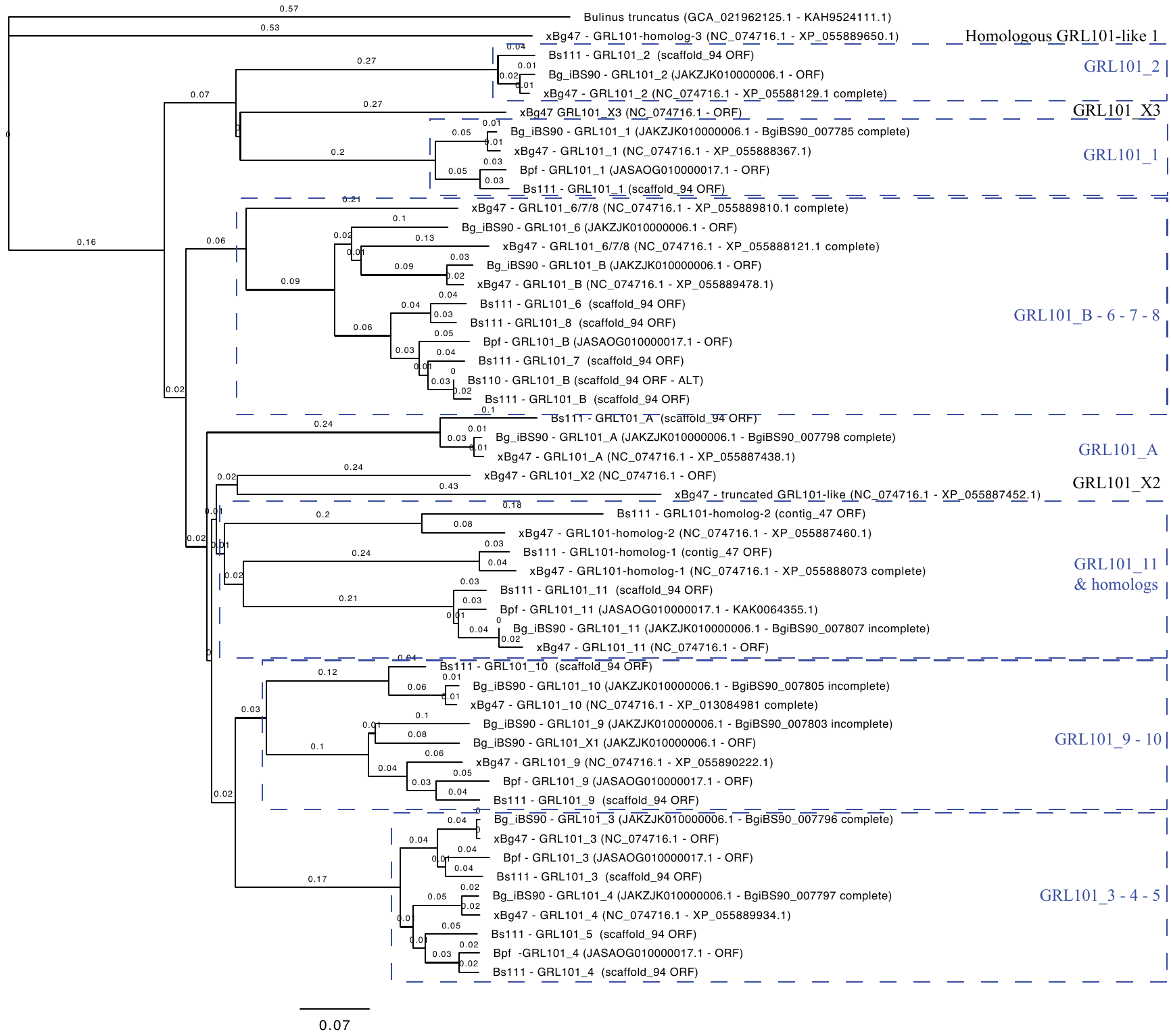

**Fig. S15** -Neighbor-Joining tree (model Jukes-Cantor) of all GRL101-like G protein coupled receptor (GPCR) protein sequences identified in *B. sudanica* (Bs111), *B. pfeifferi* (Bpf (22)) *B. glabrata* iBs90 (Bg_iBs90 (13)) and *B. glabrata* BG47 (xBioGlab47, NCBI RefSeq: GCF_947242115.1) *SudRes2* and adjacent regions. Amino acid sequenced aligned using the Geneious Alignment tool and tree constructed using the Geneious tree builder. Branch length shown reflects the number of substitutions per site. The outgroup is a GRL101-GPCR orthologous protein in *Bulinus truncatus* (GenBank Accessions: KAH9524111.1 (23)). Gene xBglab47-XP_055887452.1 is truncated (no transmembrane domain region). GRL101_1 and the homologous GRL101-like genes represent those not contained within the *SudRes2* region of *B. sudanica* (sc94:1.82-2.26 Mb) or *B. glabrata* (GCF_947242115.1, 8.74-9.07 Mb). An additional alternative protein isoform is given for *B. sudanica* protein GRL101_B, that was generated from transcript alignments of inbred line Bs110 (10).

Dataset S1 (separate file). Raw and processed sequence data statistics, showing number of Illumina PE150 reads obtained from pooled-GWAS sequencing of positive and negative snails, including technical replicates, and the final number of reads retained following bioinformatic processing that were aligned to the *Biomphalaria sudanica* genome.

Dataset S2 (separate file). Resulting data from *Biomphalaria sudanica* linkage map analysis using 210 markers analyzed with OneMap v. 3.0.0 (18), and showing relative chromosomes and positions on the *Biomphalaria glabrata* genome xgBioGlab47.1 (Accession GCF_947242115.1).

Dataset S3 (separate file). Markers used in the Amplicon panel for validating pooled-GWAS variants, *a priori* gene candidates and linkage markers. Dataset includes target variant coordinates (Contig/Site) in the *B. sudanica* genome (10), the purpose of the marker inclusion (dual-variant, singleton-variant, *a priori* gene candidate, linkage map marker) and the forward and reverse primer sequences. For each marker (except for all linkage map markers that are not applicable for this analysis) results of a dominance model (Fisher's exact tests) are given for genotyped-validation snails divided between groups A and B representing snail ancestry (majority population 1 and 2, respectively), and the results of an Additive Regression model including all genotyped-validation snails.

Dataset S4 (separate file). Gene identifiers and associated information for *Biomphalaria sudanica* genes in *SudRes1* region, including orthologous gene identifiers from the *B. glabrata* iBS90 genome (13).

Dataset S5 (separate file). Gene identifiers and associated information for *Biomphalaria sudanica* genes in *SudRes2* region, including orthologous gene identifiers, or orthologous gene coordinates, of GRL101 genes identified in *B. glabrata* iBS90 (13), *B. glabrata* xgBioGlab47.1 (Accession GCF_947242115.1) and *B. pfeifferi* (22) genome assemblies.

**SI References**

1.	T. Pennance, *et al.*, A rapid diagnostic PCR assay for the detection of *Schistosoma mansoni* in their snail vectors. *J. Parasitol.* in press.

2.	Global Schistosomiasisi Alliance Snail Vectors working group, Genomic DNA extraction from freshwater snail tissues. *Resources* 1–8 (2024). Available at: www.eliminateschisto.org/resources/protocol-genomic-dna-extraction-from-freshwater-snail-tissues [Accessed 10 July 2024].

3.	M. Martin, Cutadapt removes adapter sequences from high-throughput sequencing reads. *EMBnet. J.* **17**, 10–12 (2011).

4.	A. M. Bolger, M. Lohse, B. Usadel, Trimmomatic: a flexible trimmer for Illumina sequence data. *Bioinformatics* **30**, 2114–2120 (2014).

5.	M. Berriman, *et al.*, The genome of the blood fluke *Schistosoma mansoni*. *Nature* **460**, 352–358 (2009).

6.	H. Li, R. Durbin, Fast and accurate short read alignment with Burrows–Wheeler transform. *Bioinformatics* **25**, 1754–1760 (2009).

7.	H. Li, Aligning sequence reads, clone sequences and assembly contigs with BWA-MEM. *arXiv Prepr. arXiv1303.3997* (2013).

8.	H. Li, *et al.*, The Sequence Alignment/Map format and SAMtools. *Bioinformatics* **25**, 2078–2079 (2009).

9.	A. R. Quinlan, I. M. Hall, BEDTools: a flexible suite of utilities for comparing genomic features. *Bioinformatics* **26**, 841–842 (2010).

10.	T. Pennance, J. Calvelo, et al, The genome and transcriptome of the snail *Biomphalaria sudanica s.l.*: Immune gene diversification and highly polymorphic genomic regions in an important African vector of *Schistosoma mansoni*. *BMC Genomics* **25**, 192 (2024).

11.	H. Li, A statistical framework for SNP calling, mutation discovery, association mapping and population genetical parameter estimation from sequencing data. *Bioinformatics* **27**, 2987–2993 (2011).

12.	R Core Team, R: A Language and Environment for Statistical Computing. (2018).

13.	L. Bu, *et al.*, Compatibility between snails and schistosomes: insights from new genetic resources, comparative genomics, and genetic mapping. *Commun. Biol.* **5**, 940 (2022).

14.	N. R. Campbell, S. A. Harmon, S. R. Narum, Genotyping-in-Thousands by sequencing (GT-seq): A cost effective SNP genotyping method based on custom amplicon sequencing. *Mol. Ecol. Resour.* **15**, 855–867 (2015).

15.	D. H. Alexander, J. Novembre, K. Lange, Fast model-based estimation of ancestry in unrelated individuals. *Genome Res.* **19**, 1655–1664 (2009).

16.	J. M. Spaan, *et al.*, Multi-strain compatibility polymorphism between a parasite and its snail host, a neglected vector of schistosomiasis in Africa. *Curr. Res. Parasitol. Vector-Borne Dis.* **3**, 100120 (2023).

17.	J. Olson, *et al.*, A PCR-based diagnostic to detect reproduction between snails that vector schistosomiasis: at tool for genetic mapping studies. in prep.

18.	G. R. A. Margarido, A. P. Souza, A. A. F. Garcia, OneMap: software for genetic mapping in outcrossing species. *Hereditas* **144**, 78–79 (2007).

19.	F. Cabanettes, C. Klopp, D-GENIES: dot plot large genomes in an interactive, efficient and simple way. *PeerJ* **6**, e4958 (2018).

20.	T. Paysan-Lafosse, *et al.*, InterPro in 2022. *Nucleic Acids Res.* **51**, D418–D427 (2023).

21.	J. Hallgren, *et al.*, DeepTMHMM predicts alpha and beta transmembrane proteins using deep neural networks. *bioRxiv* 2022.04.08.487609 (2022).

22.	L. Bu, *et al.*, A genome sequence for *Biomphalaria pfeifferi*, the major vector snail for the human-infecting parasite *Schistosoma mansoni*. *PLoS Negl. Trop. Dis.* **17**, e0011208 (2023).

23.	N. D. Young, *et al.*, Nuclear genome of *Bulinus truncatus*, an intermediate host of the carcinogenic human blood fluke *Schistosoma haematobium*. *Nat. Commun.* **13**, 977 (2022).
